# A 3D Amphioxus Brain Atlas Illuminates the Blueprint of the Ancestral Chordate Brain

**DOI:** 10.1101/2025.11.27.690639

**Authors:** Che-Yi Lin, Wen-Hsin Hsu, Mei-Yeh Jade Lu, Yi-Hua Chen, Yi-Chih Chen, Yi-Hsien Su, Shen-Ju Chou, Jr-Kai Yu

## Abstract

Comparative studies on highly complex vertebrate brains have provided insights into the evolutionary history of morphological diversity. However, a long-standing question in vertebrate brain evolution is how the fundamental blueprint emerged. To probe the cellular and molecular origins of brain architecture in the chordate lineage, we performed single-nucleus RNA-seq combined with spatial transcriptomics and generated a three-dimensional cell atlas of the basal chordate amphioxus central nervous system (CNS). This atlas reveals a tripartite organization along the anterior-posterior axis of the adult amphioxus brain, with a rostral retinal/hypothalamic region followed by a di-mesencephalon and caudally, the hindbrain and spinal cord. Notably, we show that the amphioxus brain contains cell clusters resembling the zona limitans intrathalamica and midbrain-hindbrain boundary, both of which demarcate the major brain compartments. Furthermore, expression profiling and gene ontology enrichment analyses support that the cell clusters located in the rostral part of the amphioxus forebrain correspond to vertebrate retinal primordium and hypothalamic cells. However, the absence of key telencephalic marker expression in the amphioxus anterior forebrain suggests that the telencephalon likely represents a vertebrate innovation. Together, our findings establish a spatial transcriptomic framework for the amphioxus CNS and provide a critical link for understanding the evolution of the complex vertebrate brain.

## Introduction

Despite substantial variability in brain sizes, shapes, and compartmental proportions across species, all vertebrate brains are divided into three main parts, including the forebrain (prosencephalon), midbrain (mesencephalon) and hindbrain (rhombencephalon)^1^. These three regions are patterned during early development by three signaling centers: the anterior neural ridge (ANR), the zona limitans intrathalamica (ZLI), and the isthmic organizer (IsO) located at the midbrain-hindbrain boundary (MHB)^2^ (Figure 1a). This basic organization and its corresponding genetic controls are common to all extant jawless and jawed vertebrates, suggesting that this fundamental brain layout already existed at the origin of vertebrates, more than 500 million years ago (Mya)^1,3,4^. However, it remains unresolved exactly when and how this brain blueprint evolved, particularly with regard to the vertebrate forebrain, which comprises the rostrally located secondary prosencephalon (telencephalon + hypothalamus) followed by the diencephalon (Figure 1a)

**Figure 1.**
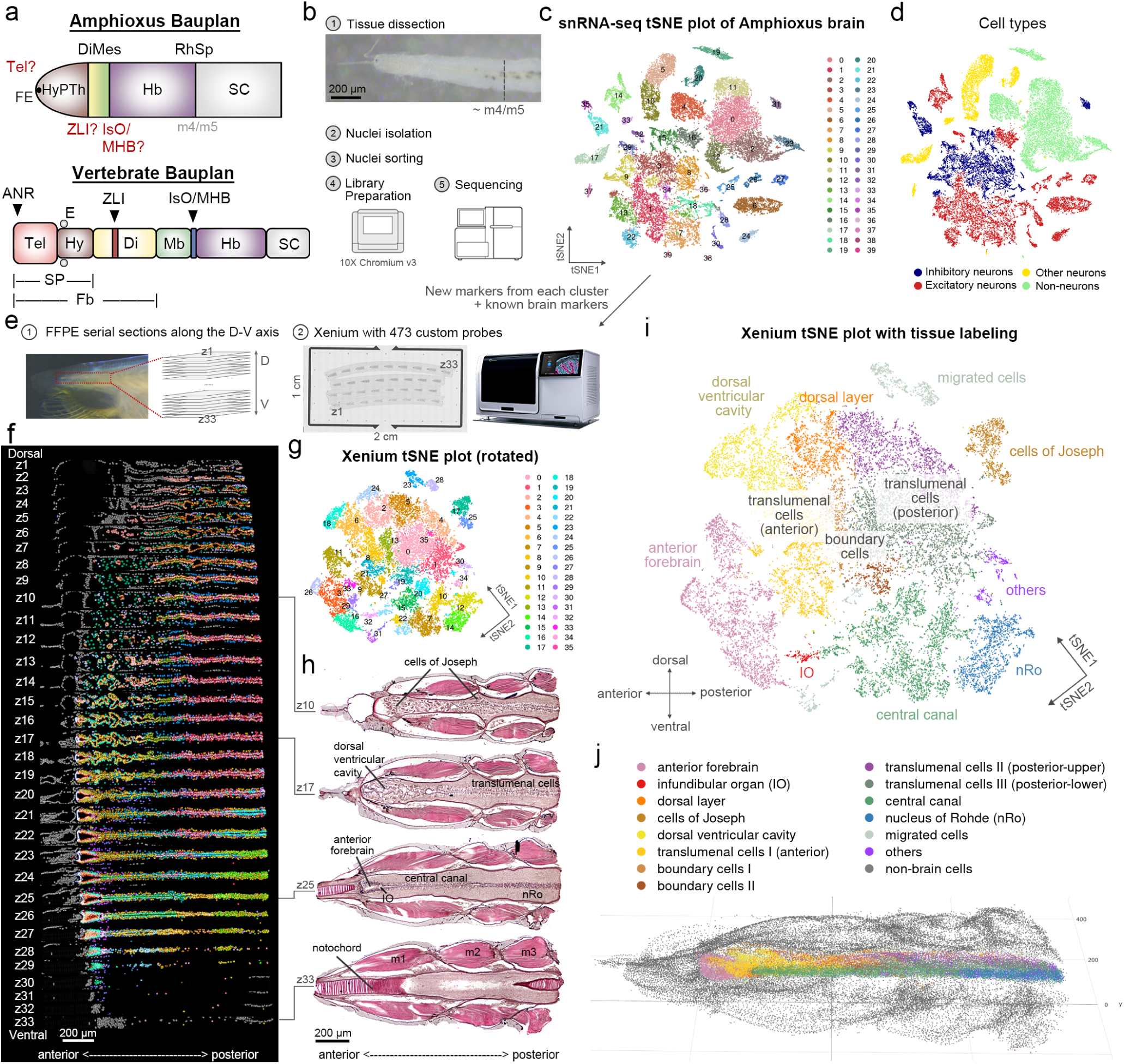
Single-nucleus and spatial transcriptomic atlas of adult amphioxus brain. **a**, Schematic representation of the Bauplan of the amphioxus and vertebrate CNS. Based on a previous study^17^, the developing amphioxus CNS can be subdivided into the hypothalamo-prethalamic primordium (HyPTh), which harbors the rostral frontal eye (FE); the di-mesencephalic primordium (Di-Mes); and the rhombencephalo-spinal primordium (RhSp), comprising the hindbrain (Hb) and spinal cord (SC). The boundary between the brain and spinal cord is at the junction of myomere 4 and 5 (m4/m4). The vertebrate CNS consists of the forebrain (Fb), midbrain (Mb), hindbrain (Hb), and spinal cord (SC). The vertebrate forebrain can be further divided into the telencephalon (Tel) and hypothalamus (Hy), which together constitute the secondary prosencephalon (SP), as along with the diencephalon (Di). In addition, three major vertebrate brain organizers are indicated: the anterior neural ridge (ANR), the zona limitans intrathalamica (ZLI), and the isthmic organizer (IsO) located at the midbrain-hindbrain boundary (MHB). The putative existence of the telencephalic, ZLI, and IsO/MHB homologous domains in the amphioxus brain remains controversial. E, eyes. **b**, Overview of the snRNA-seq experimental workflow. The dashed line denotes the boundary between the dissected brain and spinal cord, at approximately myomere 4-5 (m4/m5). Nuclei were isolated from pooled adult amphioxus brains, sorted based on 7-AAD staining, and subjected to library preparation using the 10x Genomics Chromium platform, followed by sequencing. **c**, t-SNE plot of brain snRNA-seq dataset; 40 distinct clusters were identified at a resolution parameter of 1. **d**, Projection of major cell types onto the t-SNE plot, annotated based on transcriptomic signatures (see Figure S4 and S5). **e**, Overview of the Xenium experimental workflow. Serial FFPE sections of the anterior region of the animal were mounted onto a single Xenium slide. The slide with a custom panel of 473 probes was processed on the Xenium platform for image-based transcript detection. **f**, Xenium images aligned across serial dorsal-ventral sections, oriented with anterior to the left. Brain regions were selectively retained and cropped (see Figure S9 for the full dataset). Cells are colored according to assigned cluster. **g**, t-SNE visualization of the brain spatial transcriptomic dataset, with rotated axes for clarity, showing 36 distinct clusters identified at a resolution parameter of 1.5. **h**, H&E-stained images of the sections, oriented with anterior to the left (see Figure S10 for the full dataset). IO, infundibular organ; nRo, nucleus of Rohde; m1-m3, myomeres 1-3. **i**, Projection of the annotated brain regions onto the t-SNE plot of the Xenium dataset. **j**, Reconstructed 3D point cloud model of the brain with color-coded anatomical regions.

This question may be best addressed by comparative analyses across vertebrates and closely related invertebrate chordates. All chordates share a general body plan, with a dorsal tubular central nervous system (CNS), notochord, and segmented somites^5–7^. Cephalochordates (commonly called amphioxus or lancelets) are the earliest branching chordate group and thus occupy a key phylogenetic position for understanding core features of the chordate body plan, including the organization of the CNS^5,8^. The CNS of amphioxus is much simpler than that in vertebrates, as it comprises a tubular structure with a slight anterior bulge (cerebral vesicle, CV) and a trunk/tail nerve cord (the “spinal cord”)^9–11^. A series of transmission electron microscopy (TEM) studies^12–14^, together with subsequent gene expression analyses^15,16^ have established that the cells of the single frontal eye at the rostral tip of the amphioxus CV are homologous to the retinal cells of the vertebrate paired, lateral eyes; moreover, it has been considered that the amphioxus CV largely corresponds to the vertebrate diencephalon and is followed caudally by a narrow domain reminiscent of the vertebrate midbrain^10,11,17–19^. A recent gene survey on amphioxus neurulae further revealed molecular regionalization in the developing CNS^17^. The anterior portion was subdivided along the anteroposterior axis into the hypothalamo-prethalamic (HyPTh) primordium, with a rostral frontal eye^13^, the di-mesencephalic (Di-Mes) primordium, and the rhombencephalo-spinal (RhSp) primordium (Figure 1a), and no telencephalic-like region was recognized. Interestingly, it was reported recently that orthologs of several marker genes of the vertebrate telencephalon are co-expressed in the anterior dorsal part of the adult amphioxus CV^20^. Along with the presence of some similar neuronal cell types, these gene expression observations led to an alternative hypothesis that the anterior dorsal part of the amphioxus cerebral vesicle may be homologous to the vertebrate telencephalon^20^. Altogether these results demonstrate a highly-patterned amphioxus brain along the anteroposterior axis. However, it remains unclear whether comparable secondary organizers are present in amphioxus brain. Moreover, uncertainty regarding the presence of a telencephalon-like region in amphioxus brain, and the lack of information on the cellular composition of the amphioxus HyPTh have both hindered our understanding on the evolutionary origins of the vertebrate brain (Figure 1a). To resolve these questions, a comprehensive understanding of amphioxus brain cell-type diversity is required.

Single-cell transcriptomics can reveal large gene sets that are co-expressed in individual cells^21^. Recent studies have applied single-cell RNA sequencing (scRNA-seq) to the brains of both larva and adult sea lampreys^22^, an early branching vertebrate lineage. Comparing these datasets with those from mice revealed deep conservation of neuronal cell types, as well as lineage-specific innovations. To further trace the cellular and molecular origins of the ancestral brain architecture within chordates, we combined single-nucleus RNA sequencing (snRNA-seq) and spatial transcriptomics to construct a comprehensive cell type atlas of the adult CNS in amphioxus. Our results reveal a tripartite organization of the amphioxus brain that parallels the organization in vertebrates, including comparable boundaries (ZLI-like and MHB-like) demarcating the major compartments along the anterior-posterior axis. Importantly, we found no evidence for a telencephalon-like region within the amphioxus forebrain, suggesting that the telencephalon, particularly the dorsal telencephalon (pallium), likely represents a vertebrate-specific innovation.

## Results

### 1. Single-nucleus transcriptomics reveals high cellular diversity of the amphioxus anterior neural tube

To investigate the cellular composition and heterogeneity of the amphioxus CNS, we isolated the entire neural tube and dissected it into anterior and posterior regions (henceforth referred to as the “brain” and “spinal cord” regions, respectively), for snRNA-seq analysis using the 10x Genomics Chromium platform (Figure 1b). The brain region encompasses the entire cerebral vesicle and the first few organs of Hess^9,23^, while the remaining portion of the neural tube is defined as the spinal cord. The boundary between these two regions is at the junction of myomere 4 and 5^9^. After quality control and data filtering, we obtained transcriptome data from a total of 36,144 and 30,997 high-quality nuclei for the brain and spinal cord, respectively. Compared to a previously published amphioxus neural tube dataset using the SPLiT-seq protocol^24^, our brain dataset included more genes and transcript counts per nucleus (Figure S1a-c), enabling more refined cell clustering and facilitating the identification of rare cell types. Using the FindClusters function of the Seurat tool with a resolution of 1, the brain cells were categorized into 40 distinct clusters, and spinal cord cells were categorized into 35 clusters (Figure 1c, and Figure S2a and S2b). These two datasets were readily distinguishable based on the expression of *Hox3* and *Hox4*, which are restricted to the spinal cord (Figure S2c and S2d)^25^.

To investigate whether the amphioxus brain harbors greater cellular diversity than the spinal cord, we integrated the two snRNA-seq datasets in low-dimensional space and examined the extent of gene expression overlap in brain- and spinal cord-derived cells (Figure S3a-c). Indeed, specific regions of the t-SNE dimensional reduction space contained only brain-derived nuclei and lacked representation from the spinal cord (Figure S3c). To further quantify tissue specificity, we computed a brain- and spinal cord-specific cell indices for each nucleus, then projected these scores onto the t-SNE map of each dataset (Figure S3d-f). This analysis revealed marked asymmetry between the two datasets. In the brain dataset, 14 clusters showed a median brain-specific index of 100, indicating that these clusters were almost entirely comprised of brain-derived cells (Figure S3e and g). In contrast, spinal cord-specific index values were more evenly distributed across clusters of the spinal cord dataset, suggesting that most cell types in the spinal cord were also represented in the brain dataset (Figure S3f and S3h).

To further annotate the amphioxus brain cell types, we examined expression of canonical markers for mature neurons (*Snap25* and *Map2*) and non-neuronal cells (*Quaking* and *EAAT*). This analysis allowed us to distinguish mature neuronal clusters from non-neuronal clusters (Figure S4). Further classification using excitatory neuron markers (*VGluT1* and *VAChT*) and inhibitory neuron markers (*Gad1* and *VIAAT*) revealed that most mature neuronal clusters corresponded to either excitatory or inhibitory neurons (Figure 1d and Figure S5a-f). Of note, a small subset of clusters (clusters 5, 10 and 17) expressed monoaminergic transporter genes (*VMATa* and *VMATb*), suggesting that the amphioxus brain contains monoaminergic neurons (Figure S5g and S5h). We then compared our amphioxus dataset with an scRNA-seq dataset of mouse brain using SAMap^26^, an algorithm that identifies homologous cell types across datasets based on gene expression similarity. This comparison revealed similarities between amphioxus neurons and mouse neuronal cell types spanning early embryonic (E11.0) to adolescent stages (P12-P60) (Figure S6). Consistently, putative amphioxus non-neuronal cells also corresponded to mouse non-neuronal populations (particularly radial glia) from various brain regions, such as the optic cup, cortical hem, roof plate and floor plate (Figure S7). Based on the SAMap cross-species comparison, a high degree of similarity in non-neuronal population was especially apparent during early development (E9.0-E12.0), suggesting that these non-neuronal cell types may represent the progenitor cell pools in the amphioxus adult brain. This interpretation is in line with previous anatomical descriptions of the neuroglial cell types in adult amphioxus^9,11,27^ and may reflect a prolonged neural developmental process in amphioxus. Further comparison with regionally annotated mouse cell types (Figure S7 and S8) showed that amphioxus adult neurons primarily map to mouse glutamatergic and GABAergic neurons in the forebrain, midbrain and hindbrain, particularly at late embryonic stages (Figure S8). This pattern of correlation suggests that these matching groups of brain cell types may represent homologous cell-type families dating back to the last common ancestor of chordates more than 550 Mya^8^. In sum, our snRNA-seq results support the transcriptional conservation of amphioxus neuronal and non-neuronal cell types, allowing us to align the identified cell types with putative vertebrate counterparts in distinct brain regions. Interestingly, certain cell types within several mouse brain regions showed no significant correspondence to amphioxus cell clusters (Figure S8). Thus, our results may reveal both evolutionary conservation and likely innovations in diverse vertebrate brain cell types.

### 2. High-resolution 3D spatial transcriptomic atlas reveals anterior-posterior and dorsal-ventral regionalization of the amphioxus brain

To add detailed spatial information to the identified cell types within the amphioxus adult brain, we performed imaging-based spatial transcriptomics on anterior CNS tissue of adult amphioxus. We used 10x Genomics Xenium platform with a custom panel of 473 genes (see Methods), incorporating both newly identified cluster-specific differentially expressed genes (DEGs) and established brain markers (Figure 1e, Supplementary Table 1). The spatial transcriptomic profiling captured a total of 51,540 cells from 33 formalin-fixed paraffin-embedded (FFPE) serial sections along the dorsoventral axis, with clear anatomical segregation between brain and non-brain tissues based on both low-dimensional space (Figure S9a and S9b) and spatial information (Figure S9c). Among the analyzed cells, 23,428 were identified as brain cells, which can be categorized into 36 transcriptionally distinct clusters (Figure 1f and g). We then generated a 3D transcriptomic model of the amphioxus brain by performing 3D tomography reconstruction with rigid registration of spatial transcriptomic sections (Figure 1j and 5a). Subsequently, we annotated the 3D transcriptomic map based on published studies and histological staining (Figure 1h and Figure S10). For example, we identified cells of Joseph in the dorsal region; these cells are presumed rhabdomeric photoreceptors (z10, Figure 1h)^16,28^. In addition, three ventricular cavities were identified^9^, including a dorsally located ventricular cavity (z17, Figure 1h), the anterior forebrain (z25, Figure 1h) corresponding to the hypothalamo-prethalamic primordium (HyPTh) in larvae^17,25^, and the ventrally dilated central canal that extends into the nerve cord (z25, Figure 1h). The infundibular organ (IO), which is likely secretory,^27,29,30^ lies between the anterior forebrain and the central canal (z25, Figure 1h). In the caudal region of the brain, we observed a dense ventral cluster of cells corresponding to the nucleus of Rohde (z25, Figure 1h)^11^. The nerve cord is primarily composed of translumenal cells, with identities of predominantly excitatory and inhibitory neurons (z17, Figure 1h)^9^. Based on these results, our spatial transcriptomic atlas recapitulated major cell types described in previous anatomical studies of the amphioxus brain. Interestingly, the spatial distribution of the cell clusters in the Xenium t-SNE space are organized into a distinct dorsal-ventral and anterior-posterior pattern (Figure 1i). We thus re-orientated the t-SNE plot with the anterior end on the left and the dorsal side at the top, reflecting the anatomical axes. We then used SAMap to align the transcriptomic profiles between our snRNA-seq and Xenium dataset (Figure S11). This analysis demonstrated that cell clusters identified from snRNA-seq closely matched those in the Xenium dataset at both the cluster and tissue levels, demonstrating the robustness and consistency of our cell type classification across the two independent platforms. Altogether, our spatial transcriptomics results revealed a clear regionally segregated pattern of cell types along both the dorsal-ventral and anterior-posterior axes of the amphioxus brain.

### 3. Transcriptional regionalization along the anterior-posterior axis reveals a tripartite brain organization in amphioxus

Previous studies proposed that the developing amphioxus brain can be divided into three major domains along the anterior-posterior axis^17,18,31^: HyPTh (hypothalamo-prethalamic primordium, marked by *Rx*), Di-Mes (diencephalon-mesencephalon, marked by *Sim*), and rhombencephalon (hindbrain, marked by *Pax2/5/8* and *Gbx*). Similarly, we found that the adult amphioxus brain could be subdivided along the anterior-posterior axis based on region-specific gene expression patterns (Figure 2a-c). For example, the *Nkx2.1* and *Rx* transcripts marked the anterior forebrain, while *Sim* was predominantly expressed in cells near the anterior central canal, a region likely corresponding to the Di-Mes domain. IO, marked by *Dbx*, is located between HyPTh and Di-Mes. Posteriorly, two amphioxus hindbrain markers, *Pax2/5/8* and *Gbx*, were expressed in translumenal neurons near the second myomere, consistent with the hindbrain territory.

**Figure 2.**
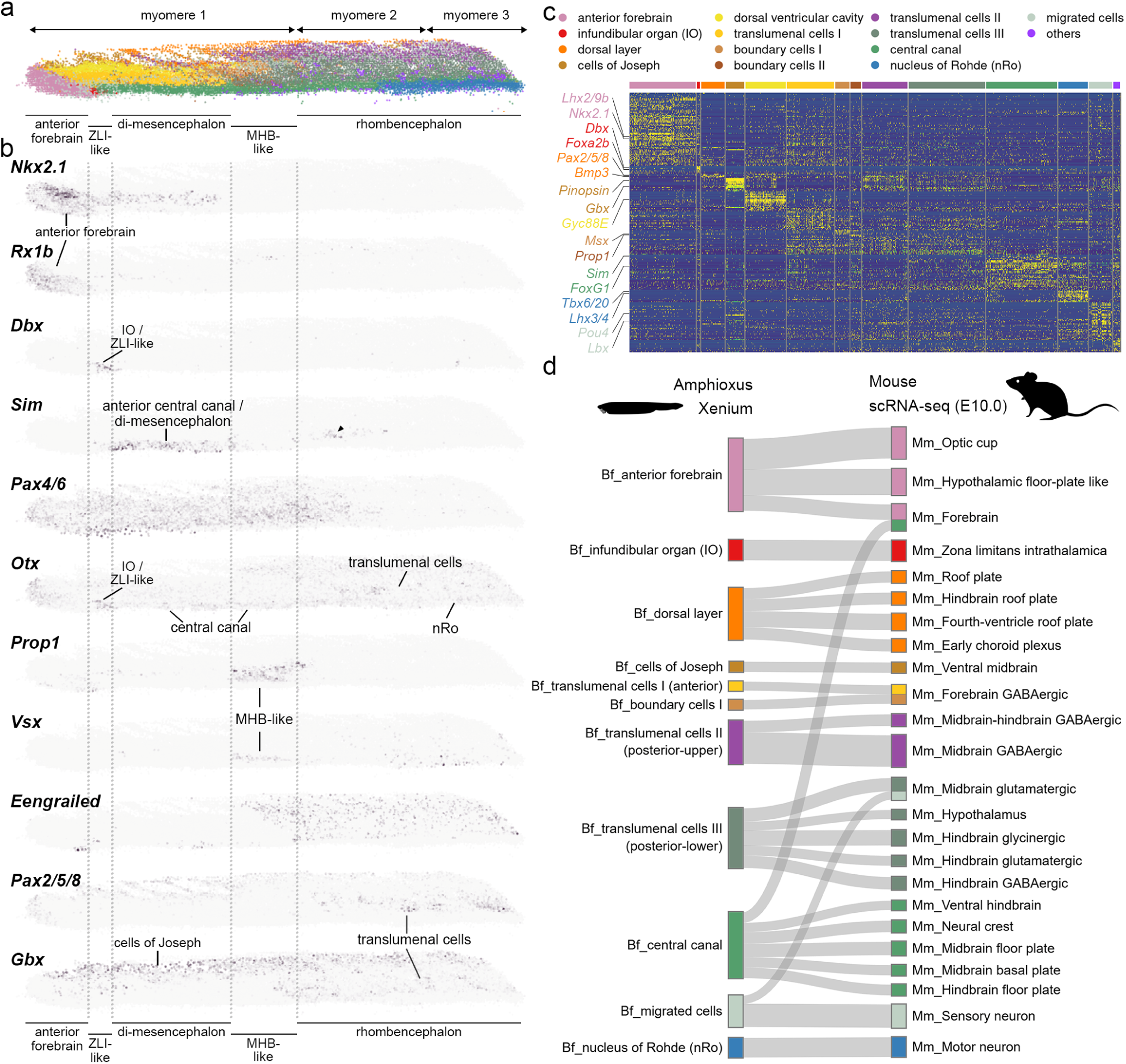
Anterior-posterior (A-P) patterning of the adult amphioxus brain. **a**, Lateral view of the 3D point cloud model of the amphioxus brain, oriented with anterior to the left and colored by the anatomical brain regions. **b**, Lateral view of the 3D point cloud model of the brain, showing expression levels and domains of individual genes. Color scales were normalized for each gene, with the darkest tone representing the maximal expression. Brain regions corresponding to major gene expression domains were labeled. **c**, Heatmap showing the top differentially expressed genes from the Xenium dataset across defined brain regions. **d**, Sankey plot showing SAMap-based cell-type mapping between the amphioxus Xenium dataset and the E10.0 mouse brain dataset. Only alignments with scores > 0.2 are displayed. Colors correspond to the amphioxus annotated brain regions. Full alignment results are shown in Figure S12.

Additional comparisons between the amphioxus and the mouse single-cell datasets^32^ revealed several corresponding cell types between the amphioxus and mouse brains (Figure 2d and Figure S12-S16). For instance, the anterior forebrain of amphioxus mapped to the optic cup, hypothalamic floor plate, and forebrain of the E10.0 mouse embryo (Figure 2d and Figure S12). In addition, anterior translumenal cells corresponded to GABAergic neurons in the mouse forebrain. In contrast, posterior translumenal cells corresponded to GABAergic and glutamatergic neurons in the mouse midbrain and hindbrain, reflecting enrichment of inhibitory and excitatory neurons in the translumenal region as previously described^9^. In addition to the general anterior-posterior correspondence between amphioxus and mouse brains, we observed a conserved dorsoventral alignment. The amphioxus dorsal layer expressed *Atoh1* (a vertebrate roof plate marker) and mapped to the mouse roof plate, while the ventrally located central canal was *Nkx2*.2- and *Foxa2-*positive and corresponded to the mouse floor plate (Figure S17), reflecting a conserved dorsoventral patterning mechanism shared between amphioxus and vertebrates^33,34^. Intriguingly, we observed a weaker tissue-level correspondence when comparing amphioxus data to later developmental stages of the mouse brain (e.g., E16, E18 and adolescent, Figure S15 and S16). Most amphioxus brain regions mapped to mature neuronal populations in the E18 mouse brain, specifically GABAergic (the most abundant inhibitory) and glutamatergic (the most abundant excitatory) neurons. These observations likely reflects the predominance of differentiated neurons and the relative scarcity of precursor cells (e.g., radial glia) at later developmental stages in mouse (Figure S6). Thus, this finding supports a temporal difference in neuronal and non-neuronal development between the two species. Taken together, our spatial transcriptomic data revealed a transcriptional regionalization along both the anterior-posterior and dorsal-ventral axis in the amphioxus brain, suggesting a conserved patterning blueprint in the chordate lineage.

### 4. Existence of a ZLI-like signaling center and the midbrain-hindbrain boundary in the amphioxus brain

The vertebrate ZLI organizer is located in the diencephalon at the boundary of the prethalamus (P3) and thalamus (P2)^35^. This region plays a crucial role in patterning the vertebrate brain, and similar gene expression patterns in the ZLI and the junction between the proboscis and collar in hemichordates led to a hypothesis that a ZLI-like signaling center was present in the common ancestor of deuterostomes^36^ but lost in the amphioxus lineage^17,36^. Here, we found expression of markers typically associated with the vertebrate ZLI (e.g., *Dbx*, *Otx* and *Foxa2b*) in the amphioxus IO (Figure 2b-c and 3a), which corresponds to cluster 38 of our snRNA-seq dataset and is marked by the expression of *Sspo*^27,29,30^ (Figure 3b and c). Notably, amphioxus ortholog of the mouse canonical ZLI signaling molecule *Wnt8*^32^ was also specifically expressed in this cluster (Figure 3c), showing strong co-expression with *Sspo* in IO cells (Figure 3d). Furthermore, other genes typically expressed in regions surrounding the ZLI (e.g., *Dlx* and *Pax4/6*) are expressed near the IO (Figure 3a). However, amphioxus orthologs of genes encoding ventral ZLI signaling components, such as *Shh* and its receptor *Ptch*, were rarely detected in the IO (Figure 3a and c).

**Figure 3.**
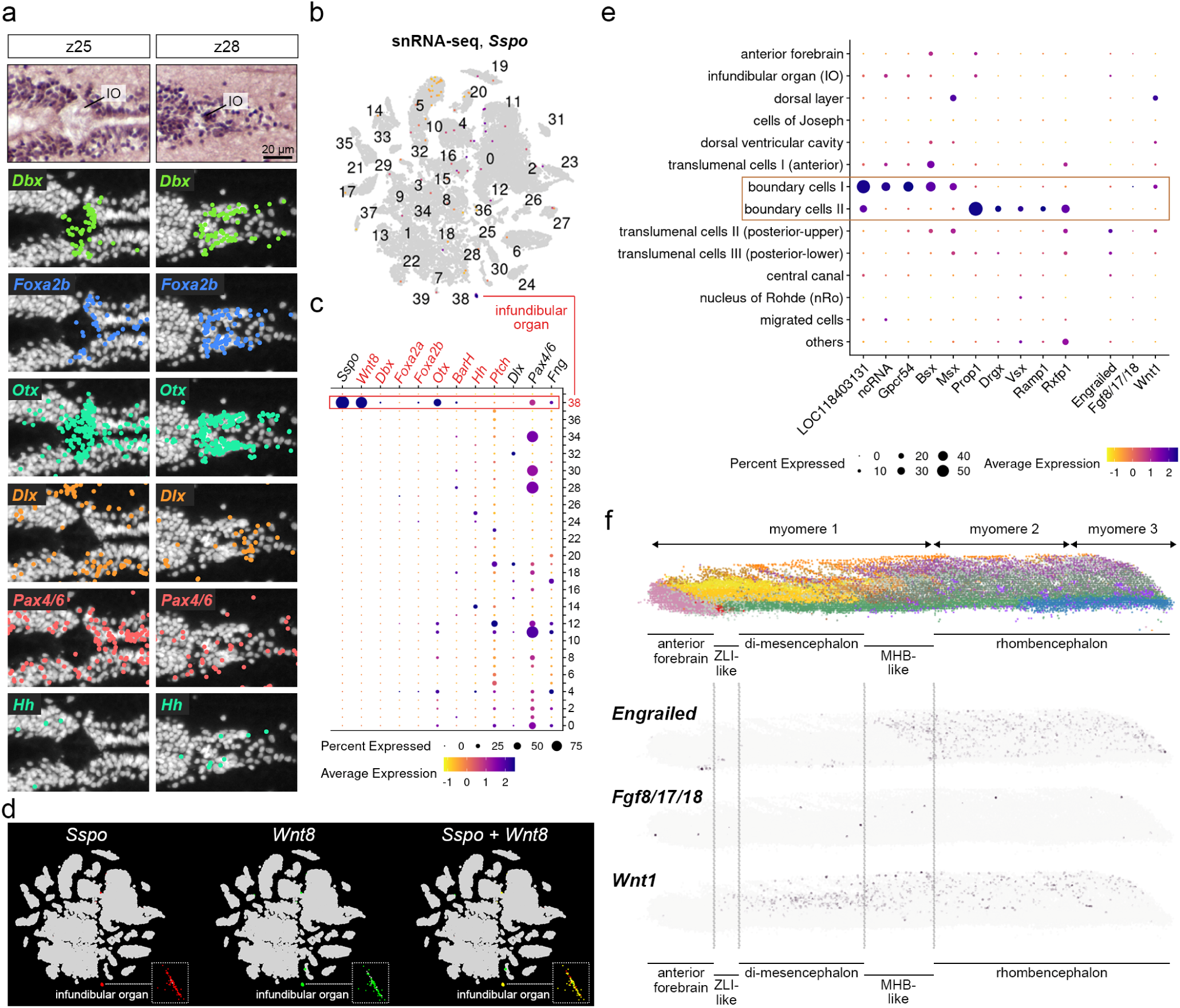
Expression of the ZLI and MHB markers in the adult amphioxus brain. **a**, H&E-stained and DAPI-stained images with Xenium spatial transcriptomic data shows corresponding localizations of ZLI-related genes around the IO. Each dot represents a single detected transcript. **b**, Feature plot of the snRNA-seq dataset showing *Sspo* expression predominantly in the IO (cluster 38). **c**, Dot plot of the snRNA-seq dataset showing the expression of ZLI-related genes across clusters. **d**, Feature plots of the snRNA-seq dataset showing co-expression of *Sspo* and *Wnt8* in IO cells. **e**, Dot plot of the Xenium dataset showing the expression of amphioxus genes expressed in the boundary cell types and amphioxus orthologs of the canonical vertebrate MHB-related genes across tissues. **f**, Transcript levels and distributions of the amphioxus orthologs of the vertebrate IsO/MHB markers, including *Engrailed*, *Fgf8/17/18* and *Wnt1*.

To further examine the possible evolutionary relationship between the amphioxus IO and the vertebrate ZLI, we further compared transcriptomes of the IO and mouse brain cell types. SAMap analysis revealed that the amphioxus IO corresponds to three mouse structures: the ZLI, the cortical hem (an embryonic signaling center that organizes hippocampal and neocortical development^37,38^), and the subcommissural organ (where *Sspo* is also expressed) (Figure S18). Notably, both the cortical hem and the ZLI are forebrain signaling centers, with cortical hem located dorsally and the ZLI positioned ventrally^39,40^. Together, our results suggest that the amphioxus secretory IO may function as a signaling center with a ZLI-like property that delineates the boundary between the HyPTh and the Di-Mes region. However, the vertebrate ZLI lies between the anteriorly located P3 and the posteriorly located P2 within the diencephalon, while the amphioxus IO is anterior to the region with expression of P3 markers, such as *Wnt7* (Figure S19). Consistently, the *Fezf*-*Irx* expression boundary that marks P3/P2 boundary in vertebrates was detected posterior to the IO, suggesting that the amphioxus IO demarcates the anterior forebrain and the Di-Mes region, rather than partitioning the P3 and P2 domains within the diencephalon as in vertebrates (Figure S19). Thus, these results suggest that the ZLI-like organizer may have slightly changed its relative position during evolution, likely coincident with the split of vertebrate P2 and P3 regions^40^ from an ancestral Di-Mes domain similar to that in amphioxus brain (Figure 5b).

The vertebrate IsO is located at the MHB and is characterized by the expression of *Fgf8/17/18* and *Wnt1* and the abutting expression domains of transcription factor genes *Otx*, *Gbx*, *Engrailed* and *Pax2/5/8* around MHB^31,41^. In our spatial transcriptomic dataset, hindbrain markers, such as *Pax2/5/8* and *Gbx*, were expressed in the translumenal cells starting at the level of the second myomere (Figure 2b). This putative hindbrain region is clearly distinguishable from the *Sim*^+^ Di-Mes region, suggesting the presence of a molecular boundary between the amphioxus Di-Mes region and the hindbrain. To further characterize this boundary, we focused on two distinct cell clusters near the caudal end of the first myomere (“boundary cells” in Figure 5a); these cells express *Prop1* and *Vsx*, serving to delineate a MHB-like region (Figure 2b). In contrast to the vertebrate MHB, expression of amphioxus *Fgf8/17/18*, *Wnt1* and *Engrailed* are not highly enriched in this boundary region (Figure 3e and f). As such, amphioxus appears to share with vertebrates only part of the molecular attributes of an MHB-like boundary between the Di-Mes and the hindbrain regions. This observation is consistent with a previously proposed evolutionary scenario^41,42^ in which the common ancestor of amphioxus and vertebrate already possessed a molecular boundary abutting the midbrain and hindbrain, and subsequently the mechanisms underlying this boundary formation have undergone partial modifications between the two lineages. During early vertebrate evolution, this ancestral MHB-like boundary may have co-opted additional regulatory genes that confer signaling center properties to this brain region, enabling further elaboration of the vertebrate brain.

### 5. Weak expression of vertebrate dorsal telencephalon (pallium) marker genes in amphioxus

The anterior part of the amphioxus brain has been suggested to be homologous to the hypothalamus and prethalamus primordium in vertebrates^17^. Moreover, gene expression of *FoxG1* during amphioxus development suggested a neuronal population similar to that in the vertebrate telencephalon^43^. It was also reported that co-expression of selected vertebrate forebrain and telencephalic markers, such as *Emx*, *FoxG1*, *Pax4/6, Nkx2.1* and *Lhx2/9,* can be detected in the dorsomedial portion of the anterior forebrain of adult amphioxus, suggesting a telencephalon-like identity^20^. However, these studies primarily relied on a limited set of patterning genes, making it difficult to distinguish true homology from potential co-option or convergent evolution.

In the developing mouse brain, *Lhx2/9* and *Pax4/6* are markers of neural progenitors and are highly expressed in both the dorsal telencephalon (pallium) and the subpallium, as well as in prethalamus (P3 domain of the diencephalon, Figure S20). Our 3D spatial transcriptomic dataset of the amphioxus brain showed transcripts of *Lhx2/9* and *Pax4/6* in the dorsomedial region of the anterior forebrain (medial, inner cells in the dorsal sections z19 and z21 in Figure 4a-b and Figure S20), consistent with the previous study^20^. In addition, this dorsomedial region exhibited weak expression of the mature neuronal marker *Snap25* (z19, Figure 4b), suggesting an undifferentiated, progenitor-like property. However, we found that *Nkx2.1* was expressed in a dorsolateral region (lateral, outer cells in the dorsal sections z19 and z21 in Figure 4a-b and Figure S20) adjacent to the *Lhx2/9* + *Pax4/6* domain. Additionally, *FoxG1*, a key pallial determinant, was only sparsely expressed in the mid-to-ventral sections (z23-z29) of the anterior forebrain (Figure 4b and Figure S20). Robust expression of *FoxG1* was confined to cells near the central canal (Figure 4a), but it was not observed in the dorsal part of the anterior forebrain. In contrast with a previous report^20^, we found that transcripts of canonical markers of the vertebrate dorsal telencephalon (pallium), including two *Emx* genes and *Tbr1*^44^, were expressed at extremely low levels across the entire brain in both the Xenium and the snRNA-seq datasets (Figure 4a and Figure S20). To further validate this observation, we searched the T1 stage single-cell RNA-seq dataset^45^ and found that transcripts of *Tbr* and *Emx* genes were also rarely detected in developing amphioxus CNS; however, *FoxG1* expression can be detected in certain cells along the HyPTh region, particularly in the middle of the HyPTh (Figure S21). This analysis provides parallel evidence to confirm that the amphioxus brain lacks expression of *Tbr* and *Emx* genes, suggesting that the dorsomedial region of the amphioxus anterior forebrain does not correspond to the vertebrate pallium, but is a progenitor-like hypothalamic cell population (based on the expression of *Lhx2/9*, *Pax4/6* and *Sox2*; Figure 4b). Taken together, these findings support the view that the telencephalon, particularly the pallium, likely represents a vertebrate innovation^5,17,46^.

**Figure 4.**
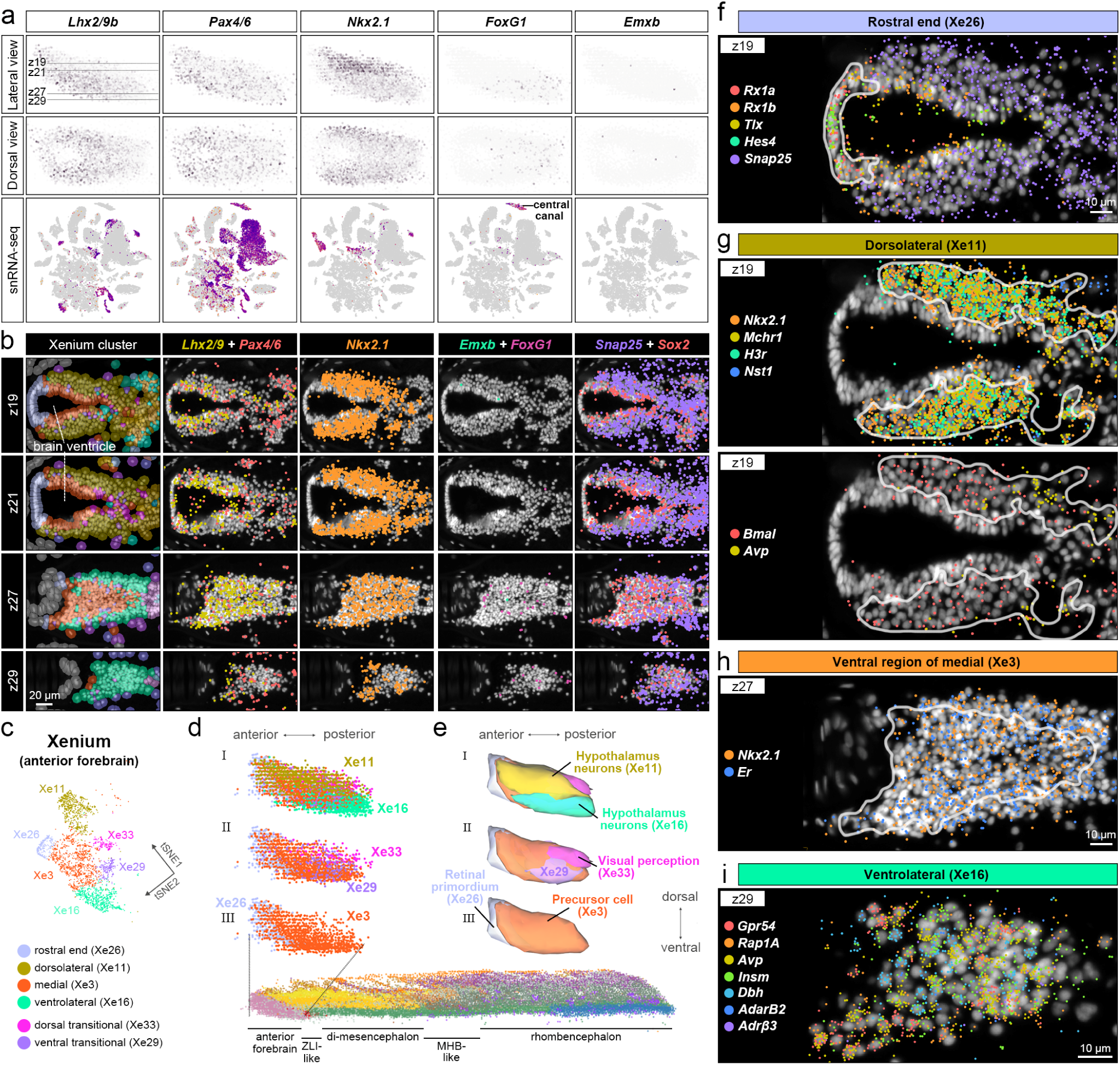
Anterior forebrain of the adult amphioxus brain. **a**, Projection of gene expression levels and distributions for anterior forebrain markers onto the 3D point cloud model (top panels for lateral view and middle panels for dorsal view) and the t-SNE space of the snRNA-seq dataset (bottom panels). **b**, DAPI-stained images with Xenium spatial transcriptomic data; shows transcript localization and cluster information around the anterior forebrain for forebrain markers, neuronal marker *Snap25*, and progenitor marker *Sox2*. **c**, t-SNE plot of the anterior forebrain subset from the Xenium dataset. **d**, Lateral view of the 3D point cloud model of the anterior forebrain, oriented with anterior to the left and colored by individual cell clusters. I, all six clusters are displayed. II, outer clusters (Xe11 and Xe16) are omitted. III, only the inner clusters (Xe26 and Xe3) are shown. **e**, Lateral view of the 3D mesh models of the anterior forebrain. **f-i**, DAPI-stained images with Xenium spatial transcriptomic dataset showing transcript localization around the anterior forebrain for markers of the rostral end (**f**), dorsolateral (**g**), ventromedial (**h**), and ventrolateral (**i**) regions.

### 6. Cell types in the anterior forebrain of adult amphioxus

To further investigate the potential functions of the amphioxus anterior forebrain and evaluate its possible homology with vertebrate brain regions, we characterized the cell types associated with its distinct regional domains. Analysis of the Xenium dataset revealed six transcriptionally distinct clusters within the anterior forebrain (Figure 1g, 1i and Figure 4c). These clusters corresponded closely with those identified in the snRNA-seq dataset, exhibiting either one-to-one or one-to-two mapping relationships (Figure S22a). Topologically, the amphioxus anterior forebrain is an ovoid structure (Figure 4d and e), with cells arranged around the ventricle cavity (Figure 4b). Leveraging spatially resolved transcriptomic data, we mapped the six anterior forebrain clusters to distinct spatial domains (Figure 4c-e and Figure S22): (1) the rostral end (Xenium cluster 26, Xe 26)), (2) the medial domain (Xe 3, cells surrounding the anterior forebrain ventricle), (3) the dorsolateral domain (Xe 11, dorsally located outer cells), (4) the ventrolateral domain (Xe 16, ventrally located outer cells), and (5-6) two transitional clusters, Xe 33 and Xe 29, were positioned between the lateral and medial domains.

We further annotated these six cell clusters according to both anatomy and gene expression profiles with GO enrichment analysis. At the rostral end (Xe 26), where the frontal eye is located^13^, transcripts of genes involved in eye development were enriched (Figure 4f), the identified genes included *Tlx*/*NR2E1* and *Hes* genes, which are known to maintain retinal stem cell identity^47–49^, as well as *Rx*, a key regulator of retinal stem cell proliferation^50^. The absence of expression of the mature neuronal marker *Snap25* in this region suggests a developmental state resembling the vertebrate retinal primordium, prior to neuronal differentiation. GO analysis revealed that many DEGs in this region are associated with negative regulation of neurogenesis and neuron differentiation, as well as camera-type eye development (Figure S23a). The medial domain of the anterior forebrain (Xe 3) expressed *Sox2*, *Lhx2/9* and *Pax4/6* (z19 and z21, Figure 4b), suggesting that this inner region may correspond to the vertebrate brain ventricular zone and retains neurogenic potential. GO enrichment analysis further supported the idea that this domain contains neuronal precursor cells (Figure S23b). Notably, the ventral region, which constitutes the ventricular floor, also expressed mature neuronal markers, including *Snap25* (Figure 4b, z27) and the estrogen receptor *Er* (Figure 4h), indicating a heterogeneous cellular composition.

The dorsolateral (Xe 11) and ventrolateral domains (Xe 16) plus the two transitional cell clusters (Xe 33 and Xe 29) exhibited robust expression of *Snap25* (Figure 4b), indicative of mature neurons. The dorsolateral domain (Xe 11) showed strong expression of *Mchr1* (Figure 4g); of note, the ligand of this receptor, *Mch*, is primarily produced in the vertebrate lateral hypothalamus and is typically involved in feeding and energy homeostasis^51^. We also observed expression of circadian-associated genes such as *Bmal* and *Avp*^52^ in this cell cluster (Figure 4g). Consistently, we found that DEGs in Xe 11 are enriched for GO terms associated with hypothalamic functions, including feeding behavior, locomotion, fear response and sleep regulation (Figure S23c). The ventrolateral domain (Xe 16) was marked by the expression of *Avp* and *Insm* (Figure 4i), and compared to the dorsolateral domain (Xe 11, Figure 4g), the Xe 16 cell cluster has higher molecular heterogeneity and may contain a mixture of neuronal populations that express partially overlapping genes (Figure 4i). GO enrichment analysis revealed that this region is associated with pathways related to learning, cognition, thermosensory behavior, xenobiotic metabolism, and hyperosmotic salinity response (Figure S23d). These data suggest that the combined dorsolateral (Xe 11) and ventrolateral (Xe 16) domains of the amphioxus forebrain (Figure 4e) likely correspond to the vertebrate hypothalamus. Interestingly, we noticed that the ventrolateral domain (Xe 16) might correspond to a region of the amphioxus larva brain that previously was annotated as HyPTh posterior pituitary (neurohypophysis-like neurons), based on the expression of *Insm* (Figure 4i), *Lhx3/4*, and *Prop1* (Figure S20)^45^, although this region lacks expression of the pituitary marker *Pit-1*^45^.

We found that the dorsal transitional cell cluster (Xe 33) is likely associated with photosensitivity, as several opsin-related genes were enriched in this region (Figure S22c). In further support of this idea, GO enrichment analysis showed that this cell cluster is associated with light detection and visual perception (Figure S23e). In contrast to the rostral end cell cluster (Xe 26), which exhibited a retinal precursor cell-like state, Xe 33 is likely a mature neuron population with photoreceptive function. This observation of multiple cell clusters with visual function is not unexpected, as photoreceptive capacity has been proposed for cells in multiple regions of the nervous system in amphioxus larvae, including the frontal eye, lamellar body, Joseph cells and organs of Hesse^16,53–55^. Notably, some of these cells altered their morphology during metamorphosis. For example, the lamellar body disappears during metamorphosis, while the remnants of individual lamellar cells protrude into the dorsal ventricular cavity in adults^9,56^. Finally, we did not find uniquely enriched GO terms in the ventral transitional cell cluster (Xe 29, Figure S23f), suggesting that this region has an expression profile that overlaps with the lateral and medial domains.

Taken together, our results suggest that the cell populations in the rostral end (Xe 26) and the medial domain (Xe 3) of the amphioxus forebrain may respectively correspond to the evolutionary precursors of the vertebrate retinal primordium and the non-pallial forebrain ventricular zone (Figure 4e). The remaining regions of the anterior forebrain, the dorsolateral (Xe 11) and ventrolateral (Xe 16) domains (Figure 4e), consist of mature neurons with molecular features similar to those in the vertebrate hypothalamus. In addition, the relatively high molecular heterogeneity within the ventrolateral domain (Xe 16) raises the possibility that individual neurons in this domain may have distinct functions. If we assume this pattern corresponds to the chordate ancestral condition, it would be likely that further segregation of cell types and the clear subdivisions of the hypothalamus may have emerged during early vertebrate evolution.

## Discussion

In this study, we reconstructed a 3D spatial transcriptomic atlas of the amphioxus central nervous system, enabling an unprecedented single-cell-resolution view of its molecular architecture. This spatially-resolved cell type atlas of the adult amphioxus brain provides a critical phylogenetic reference for tracing the evolutionary origin of the chordate brain, which can be used to infer how the complex vertebrate brain evolved. By integrating Xenium spatial gene localization data and performing cross-species transcriptomic comparisons with vertebrate brains, our study bridges a long-standing gap between developmental gene expression and adult brain organization, thereby offering a comprehensive resource for comparative neurobiology and evolutionary developmental biology.

Compared with previous gene expression studies that primarily inferred amphioxus brain regionalization from a limited set of molecular markers, our analysis reveals the transcriptional organization along the anterior-posterior and dorsal-ventral axes of amphioxus brain. Our data suggest a neuroanatomical blueprint that mirrors vertebrate brain organization yet has some key differences. We show that along the anterior-posterior axis, the adult amphioxus brain can be divided into three major segments based on transcriptomic profiles: the anterior forebrain (comprising a retinal primordium and hypothalamus), the Di-Mes, and the hindbrain (Figure 5). These segments are consistent with regions previously identified from gene expression patterns during development^17^. In addition, the three brain domains are bordered by spatially restricted cell clusters. One comprises the cell populations within the amphioxus IO that co-express ZLI-related genes, such as *Wnt8*, *Dbx*, *Foxa2b* and *Otx*. The other consists of posteriorly located MHB-like boundary cells that express homeodomain transcription factors such as *Prop1* and *Vsx* (Figure 2 and Figure 5). These findings challenge the previous assumption that neither a ZLI nor an MHB exists in amphioxus^36^. Importantly, we could not detect expression of canonical MHB signaling genes, such as *fgf8/17/18* and *Wnt1*, within this MHB-like brain region. It is therefore possible that instead of being patterned by local signaling interactions, the MHB-like brain boundary in amphioxus could be determined by cross-inhibitory regulation among transcription factors (e.g., *Prop1*, *Vsx* and other homeodomain TFs, see Figure 3); the expression domains of these transcription factors are likely under the control of the overall AP patterning mechanism along the entire body. During early vertebrate evolution, this ancestral MHB-like boundary may have co-opted additional regulatory genes that confer signaling center properties to this brain region, enabling further elaboration of the vertebrate brain. However, an MHB-like domain appears to be present at the collar-trunk coelom of the hemichordate *Saccoglossus kowalevskii*, and this MHB-like domain expresses the canonical MHB signaling genes *fgf8/17/18* and *Wnt1*^36^. Thus, it remains possible that the amphioxus MHB-like region could represent a secondary modification from the ancestral condition. Still, it should be noted that in *S. kowalevskii*, the reported expression patterns of several key TF genes around its MHB-like domain are not congruent with those in the vertebrate MHB region^36^. Thus, the ancestral genetic regulatory mechanisms patterning this boundary area remain unsettled. Nevertheless, our results reveal a distinct boundary between the amphioxus Di-Mes and the hindbrain regions, which corresponds to the vertebrate MHB and is potentially similar to the collar-trunk boundary in hemichordates. Interestingly, this amphioxus boundary appears to divide the CNS into a sensory neuron-dominant rostral region (anterior forebrain + Di-Mes) and a motor neuron-dominant caudal region (hindbrain + spinal cord), suggesting the regionalization may be inherited from an ancient subdivision of the nervous system in the chordate common ancestor.

**Figure 5.**
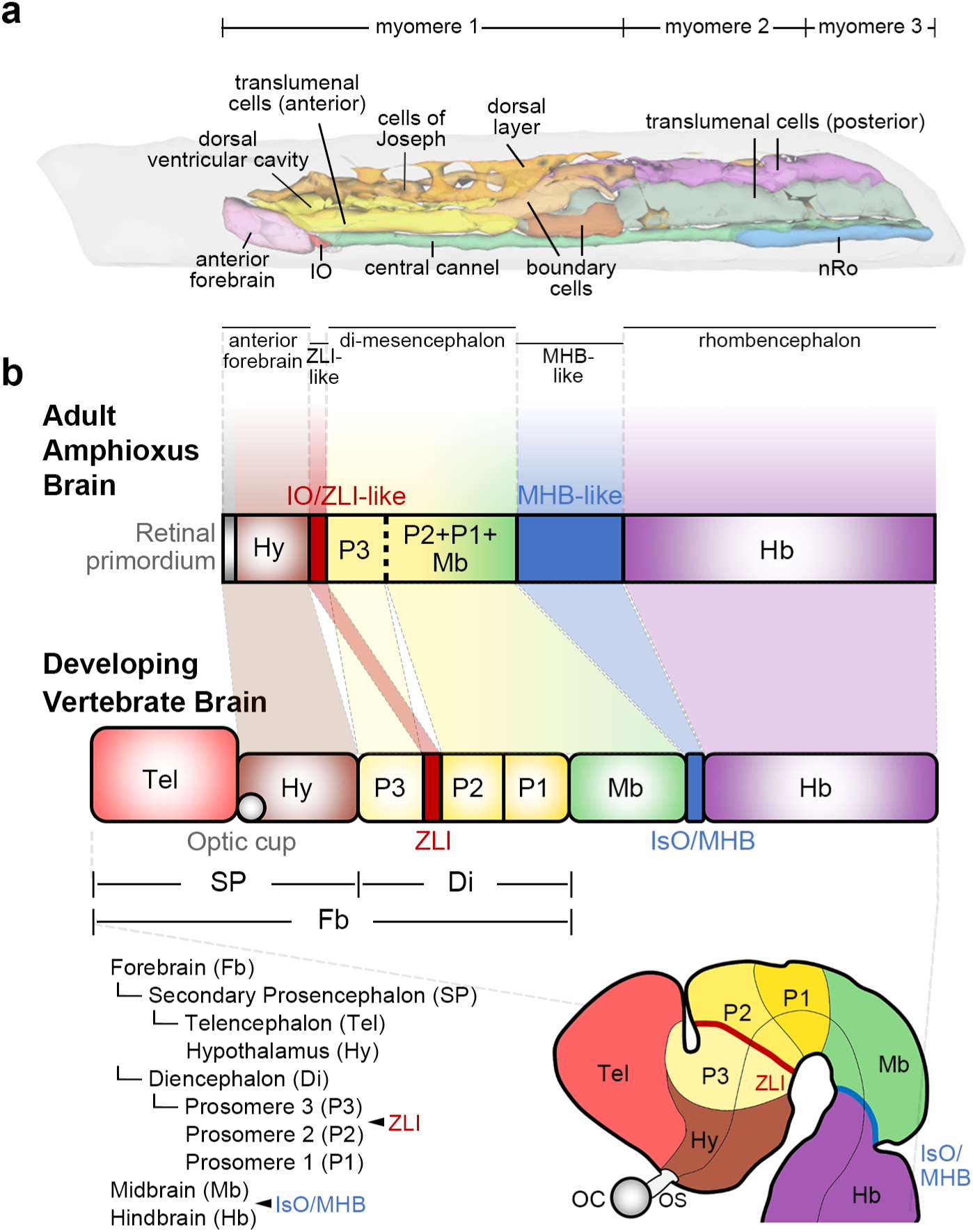
Anteroposterior organization of amphioxus and vertebrate brains. **a,** 3D mesh models of the adult amphioxus brain rendered with optimized lighting and transparency settings. Non-brain tissues are shown in gray, and brain regions are labeled in accordance with Fig. 1j. **b**, A schematic summary of the anteroposterior organization of the adult amphioxus brain and the developing mouse brain. Based on our integrative transcriptomic and spatial analyses, the adult amphioxus brain can be subdivided into three principal domains: (1) the anterior forebrain, corresponding to the vertebrate retinal primordium and hypothalamus; (2) the di-mesencephalic region, corresponding to the vertebrate diencephalon and midbrain; and (3) the rhombencephalic/hindbrain region, corresponding to the vertebrate hindbrain. Distinct molecularly defined cell populations demarcate these brain compartments. Anteriorly localized cells within the amphioxus IO express ZLI-associated genes, and the posteriorly located MHB-like boundary cells were observed. OC, optic cup; OS, optic stalk. The dashed line within the amphioxus di-mesencephalic region indicates the boundary between the expression domains of *Fezf* and *Irx*.

Of note, a recent scRNA-seq study^45^ considered the region of the amphioxus larval brain expressing *sim* to be the hypothalamus, since vertebrate *Sim* genes are expressed in the hypothalamus^57^ (Figure S24a-d). However, this assignment leads to ambiguity in anatomical interpretation of a structural sequence comprising the HyPTh hypothalamus (*Sim*⁺), IO (*Sim*⁻), and Di-Mes (*Sim*⁻), rather than HyPTh hypothalamus (*Sim*⁻), IO (*Sim*⁻), and Di-Mes (*Sim*⁺) (Figure S24e and f)^17,58,59^. In our Xenium spatial transcriptomic dataset, *Sim*⁺ cells were located immediately posterior to the IO, surrounding the ventral central canal, and terminated near the end of the first myomere (Figure 2a and b). Notably, *Sim* expression was also observed in some rhombencephalon cells in both larval (Figure S24b)^45^ and adult amphioxus datasets (Figure 2b, arrowhead), indicative of direct development without major remodeling between the larval and adult stages (Figure 2b and Figure S24b). By comparison, the mouse single-cell dataset showed that *Sim* genes are also expressed in the ventral midbrain (Figure S24c and S24d)^32^, supporting the interpretation that the *Sim*⁺ cells in amphioxus central canal reside in the Di-Mes domain, rather than in the more rostral hypothalamus region (Figure S24f). Thus, our data provide critical information for pairing anatomical structure and transcriptomic data to construct a comprehensive map of the cell types in the amphioxus brain.

Our data do not support the existence of a dorsal telencephalon-like region in the amphioxus adult forebrain. In contrast, a recent report suggested that the dorsoanterior region of the adult amphioxus brain expresses vertebrate telencephalic markers^20^. In our study, no clear expression of *Emx* and *Tbr1* (two of the canonical telencephalic genes) was detected in the amphioxus anterior forebrain. Additionally, we detected only very weak expression of *FoxG1* in the ventral part of the anterior forebrain; its expression was located within the Xe16 cell cluster, which corresponds to the hypothalamus neuron population (Figure 4a-e). This discrepancy might be attributable to differences in methodology and resolution across platforms. The earlier conclusion was based primarily on conventional *in situ* hybridization, which has limited sensitivity and specificity to discriminate transcripts that are expressed at very low levels from background signals. Here, we employed two technically independent high-resolution approaches (see Methods). Our average sequencing depth exceeded 40,000 reads per nucleus, and the quality control analysis showed that the sequencing saturation slope reached a plateau near its upper bound (Figure S1d). Therefore, we interpret the near background-level expression of *Emx* and *Tbr1* to most likely reflect their absence, rather than a technical limitation. Additionally, the imaging-based Xenium platform with a dual-probe design (requiring hybridization of two probes to adjacent sites on the target RNA) and rolling-circle amplification revealed only a few cells exhibiting near background levels of *Emx* and *Tbr1* (Figure 4, and Figure S20). Together, with these two complementary datasets, our results strongly support the conclusion that orthologs of vertebrate telencephalic markers *Emx* and *Tbr1* are not expressed in the amphioxus anterior forebrain. Thus, previous chromogenic *in situ* hybridization results may have reflected methodological limitations rather than providing robust evidence for a telencephalon-like domain in amphioxus.

Intriguingly, in the anterior part of the amphioxus forebrain, we observed clear expression of *Lhx2/9*, whose vertebrate co-orthologs (*Lhx2* and *Lhx9*) are expressed in both the telencephalon and in the hypothalamus (Figure S20). The expression of amphioxus *Lhx2/9* was detected in the cells located in the medial region of the rostral forebrain (Xe 3), but expression was not observed in the rostral end (Xe 26) (Figure 3e). The lack of overlap between the *Lhx2/9* expression domain and telencephalon-specific marker genes (*Emx* and *Tbr1*) in amphioxus forebrain further precludes the existence of a region that is equivalent to the telencephalon in vertebrates. Based on the expression profiles and GO enrichment analysis (Figure S20 and S23), we propose that cells in Xe 3 cluster have neurogenic property and constitute the ventricular zone of the hypothalamus. Additionally, the rostral end, the lateral domains, and the transitional cells in the anterior forebrain of amphioxus are most likely related to vertebrate cells located in the retinal primordium, the hypothalamus, and cells involved in visual perception, respectively. We further postulate that after the two rounds of whole genome duplication (2R-WGD) events during early vertebrate evolution^60^, some of the duplicated transcription factor genes (such as *Lhx2* and *Lhx9*) may have undergone regulatory changes, enabling the co-option of particular duplicates for the development and elaboration of telencephalon in the vertebrate lineage. Further studies on the functional diversification of key regulatory genes after 2R-WGD will provide critical knowledge for understanding this evolutionary transition.

Our finding that the anterior end of the amphioxus brain contains cells corresponding to the vertebrate retinal primordium and hypothalamic cells is in line with a recent study showing that these cell fates are specified by a conserved anterior gene regulatory network (aGRN), which is shared among all deuterostome lineages during early development^61^. In non-chordate deuterostomes (echinoderms and hemichordates), this aGRN controls development of the apical organ, a sensory structure of the larvae. Thus, the amphioxus anterior brain likely represents an evolutionary intermediate linking the primitive anterior sensory organ in the deuterostome common ancestor to the internalized, elaborated vertebrate brain. Taken together, our results suggest that the ancestral chordate brain comprised the anterior forebrain (retinal primordium + hypothalamus), the Di-Mes, and the hindbrain, while the telencephalon likely represents a vertebrate innovation that evolved from the ancestral rostral forebrain region via co-option of additional regulatory genes driving its dorsal expansion.

In summary, we constructed a 3D spatial transcriptomic atlas of the amphioxus central nervous system using snRNA-seq and spatial transcriptomic technologies. We found that compared to the spinal cord, the anterior neural tube of amphioxus is transcriptionally more complex, suggesting a brain-like regionalization rather than a simple anterior extension of the neural tube. This conclusion was further supported by Xenium spatial gene expression data and cross-species transcriptomic comparison with vertebrate brains. Together, our results demonstrate that the overall anterior-posterior and dorsal-ventral axes of the amphioxus brain broadly correspond to those in vertebrates, supporting a conserved neuroanatomical blueprint which likely represents the ancestral chordate brain organization. Furthermore, we uncover a partially conserved MHB-like region with distinct molecular property in amphioxus brain, and the lack of telencephalon counterpart in the amphioxus anterior forebrain, providing crucial information to infer the evolution of the vertebrate complex brain. We believe that future studies on the functional changes of key regulatory genes after 2R-WGD will shed more light into the genetic underpinning of brain elaboration in the vertebrate lineage.

## Methods

### 1. Sample and library preparation for snRNA-seq

Adult amphioxus (*Branchiostoma floridae*), aged 1-2+ years and measuring approximately 3-4 cm in body length, were used in this study. Animals were maintained under standard laboratory conditions prior to experimentation. Brain and spinal cord tissues were manually dissected using #5 forceps under a stereomicroscope in DEPC-treated calcium- and magnesium-free artificial seawater (4°C). The dissection was performed in a Petri dish positioned on a chilled platform and cooled with cold packs. The boundary between the brain and spinal cord was defined as the junction between myomeres 4 and 5. Brain tissue from 3-4 animals and spinal cords from individual animals were collected into separate tubes and immediately snap-frozen in liquid nitrogen. Samples were stored in the vapor layer of liquid nitrogen prior to downstream processing. Single-nucleus suspensions were prepared using the Chromium Nuclei Isolation Kit (10x Genomics) with slight modifications. Briefly, approximately 70 amphioxus brains or 8 spinal cords were pooled for each reaction, to achieve a similar total nucleus yield for each tissue type. Tissue was homogenized for 2-3 min in 100 µl of lysis buffer using a pestle. An additional 400 µl of lysis buffer was then added to the homogenate, which was subsequently applied to a nuclei isolation column and centrifuged according to the manufacturer’s instructions. Of note, the debris removal buffer provided in the kit failed to effectively eliminate tissue debris, so the debris removal step was omitted. Instead, the nuclei pellet was washed directly with a wash buffer containing 1% BSA (Sigma) and 1:40 RNase inhibitor (10x Genomics) in 1x PBS (Corning). The resulting pellet was resuspended in resuspension buffer (1% BSA, 1:40 RNase inhibitor, and 1:100 7-AAD in 1x PBS) and subjected to fluorescence-activated cell sorting (FACS). For each reaction, a total of 180,000 7-AAD^+^ singlet nuclei were sorted into the collection buffer, yielding a final concentration of 1% BSA and 1:40 RNase inhibitor in 1x PBS post-sorting. The nuclei suspension was then concentrated to approximately 1,000 nuclei/μl and examined using Trypan Blue staining under an EVOS M7000 imaging system (Invitrogen). Quality of the nuclei was assessed based on membrane integrity and morphological criteria, with intact nuclei displaying contiguous membranes and minimal to no blebbing (classified as Class A or B according to 10x Genomics guidelines). The centrifugation conditions for both wash and concentration steps were 600 rcf for 7.5 min at 4°C. Approximately 16,000 nuclei were subsequently subjected to the Chromium Single Cell 3’ Gene Expression platform (10x Genomics, v3 chemistry) for single-nucleus transcriptomic profiling, with an estimated recovery rate of approximately 65%. For the brain samples, five libraries were generated from three biological replicates. Similarly, four spinal cord libraries were prepared from two biological replicates. Except for one brain sample, technical replicates were generated by loading the same nuclei suspension into two separate wells on the 10x Chromium chip. Library preparation was performed according to the manufacturer’s protocol. Sequencing was carried out on an Illumina NextSeq 2000 system using the P3 reagent kit, targeting a read depth of over 40,000 reads per nucleus. Sequencing saturation analysis via downsampling demonstrated that the sequencing depth had reached or exceeded the reasonable saturation point (Figure S1c).

### 2. Data analysis for snRNA-seq

UMI-based gene expression count matrices were generated using Cell Ranger (v7.1.0, 10x Genomics) with the ‘include introns’ mode. The amphioxus genome and gene annotation used for alignment were downloaded from NCBI (GCF_000003815.2)^60^. Ambient RNA was removed using the remove-background function from the CellBender^62^ (v0.3.0). The resulting count matrices from each sample were then processed using the Seurat^63,64^ R package (v4.3.0) for downstream analysis. A minimum gene expression threshold of 150 genes per cell was applied for quality control. Doublet cells were identified and removed using scDblFinder^65^ (v1.16.0) with default settings. Each dataset was normalized using the LogNormalize method implemented in Seurat. For data integration across samples, the Canonical Correlation Analysis (CCA) method was employed. Data visualization was performed using the Seurat, scCustomize (v3.0.1, https://doi.org/10.5281/zenodo.5706430) and SeuratExtend^66^ (v1.1.4) R packages. A tissue-specific index was computed to assess the local enrichment of brain and spinal cord cells in the two-dimensional t-SNE space. A KD-tree was constructed using the t-SNE coordinates via the scikit-learn Python library, in order to enable an efficient nearest-neighbor search. For each cell, the 20 closest neighbors were identified, and the proportion of neighbors annotated as brain or spinal cord was calculated to quantify local tissue-type enrichment. Differentially expressed genes (DEGs) were identified using the FindAllMarkers function from the Seurat R package. Mouse brain data was obtained from a previously published study^32^ and visualized using the Scanpy^67^ package in Python.

### 3. Sample preparation for Xenium

Serial FFPE sections were prepared. After removing the epidermis, the anterior region containing the first three myomeres of adult amphioxus was dissected and immediately fixed in 10% neutral buffered formalin (Sigma) for 20-24 h at 4°C. The adjacent region containing the fourth to sixth myomeres was concurrently fixed under the same conditions to serve as an RNA quality control (QC; e.g., DV200 analysis). After fixation, samples were subjected to dehydration through sequential ethanol treatments (70%, 75%, 80%, 85%, 90%, and three changes of 100% ethanol, each for 30 min). During the final 100% ethanol step, tissues below the notochord, including oral tentacles, were trimmed to facilitate orientation in embedding molds. Samples were then cleared using HistoClear II (National Diagnostics), initially in a 1:1 ethanol:HistoClear II solution for 20 min, followed by two treatments in pure HistoClear II for 30 min each. Cleared samples were then infiltrated with paraffin wax (Leica) through three successive 30-min immersions before final embedding. Embedded tissue blocks were stored at 4°C until further processing.

Before sectioning tissues onto Xenium slides, the tissue region (myomeres 4-6) for QC was sectioned. The QC sections were collected into tubes for RNA extraction and DV200 analysis. RNA was extracted using the RNeasy FFPE Kit (Qiagen). A minor modification of performing DNase I digestion directly on-column was implemented to improve DNA removal efficiency. The integrity of the extracted RNA was assessed via a High Sensitivity RNA Analysis Kit on a Fragment Analyzer (Agilent), and samples with more than 80% of RNA fragments exceeding 200 bp were selected. Finally, the corresponding tissue blocks containing the anterior region (myomeres 1-3, including brain structures) were serially sectioned at 5 μm thickness using a Leica microtome. A complete set of brain sections from one individual animal was mounted onto a single Xenium slide. The slides were then incubated at 42°C for 3 h on a hot plate (Leica), inspected using an EVOS M7000 microscope (Thermo Fisher Scientific), and stored in a desiccator until further processing.

### 4. Xenium experiment

The Xenium experiment was performed according to the manufacturer’s protocols (CG000580, CG000582 and CG000584, 10x Genomics). In brief, deparaffinization was carried out on Xenium slides containing FFPE tissue sections. The slides were incubated at 60°C for 2 h to melt the paraffin, followed by two sequential immersions in xylene to remove residual wax. Rehydration was performed through a graded ethanol series (100%, 96% and 70%), ending with a final rinse in nuclease-free water. After deparaffinization, decrosslinking was carried out using the Decrosslinking Buffer provided in the Xenium kit.

Xenium In Situ Gene Expression assays were then performed according to the manufacturer’s protocol with the following steps. Probe hybridization was conducted at 50°C overnight to allow specific binding of RNA-targeting probes. Probe ligation was performed using the Ligation Mix at 37°C for 2 h to circularize hybridized probes. Rolling circle amplification was carried out at 30°C for 2 h to generate a robust and strong signal. Autofluorescence quenching and nuclei staining were performed as described in the protocol to reduce background signal and facilitate nuclear visualization. Prepared slides were subsequently processed on the Xenium Analyzer for sequential on-instrument imaging cycles and transcript decoding. The outputs included DAPI images overlaid with cell clusters and transcript locations, as well as transcript count matrices for each detected cell. Cell segmentation was performed based on DAPI staining, using nuclear boundary expansion with a 5-μm radius.

Following the Xenium experiment, H&E staining was performed according to the manufacturer’s protocol (10x Genomics). Stained slides were imaged using a ZEISS microscope equipped with a 20× objective lens and the TissueFAXS imaging system. QuPath (v0.5.1) was used for file format conversion. Image stitching and background removal were performed using Adobe Photoshop (v2025).

### 5. Data analysis for Xenium

Transcript count matrices from the Xenium dataset were processed using the Seurat R package (v4.3.0) for downstream analysis. The data were normalized using the LogNormalize method. Standard Seurat workflows were then applied, including data scaling, linear dimensionality reduction (PCA), clustering, and non-linear dimensionality reduction (UMAP and t-SNE). Cells corresponding to brain regions were subsequently subsetted and reprocessed using the same pipeline to obtain brain-specific clustering results. The resulting t-SNE plot of brain cells was manually rotated with a custom code to match the anatomical orientation. Clustering information was imported into Xenium Explorer (v3.2.0), where DAPI images were visualized in combination with either clustering results or transcript locations. Differentially expressed genes were identified using the FindAllMarkers function from the Seurat R package. Gene Ontology (GO) annotation for amphioxus was performed using the Blast2GO pipeline within OmicsBox^68–70^; BLAST searches were restricted to the metazoan subset. GO enrichment analysis was carried out using the clusterProfiler^71^ R package (v4.10.1).

### 6. 3D model reconstruction

The 2D coordinates of individual nuclei were retrieved from the Xenium output. In the original Xenium coordinate system, the Y-axis increases downward (negative Y-values point upward). To conform to standard spatial orientation, the Y-axis was first inverted so that positive Y-values point upward. Subsequently, each 2D section was then translated so it was centered at the origin (0,0) in the XY plane. To achieve 3D reconstruction, sequential pairwise rigid registration was performed across sections. Instead of aligning all sections to a single reference (e.g., the first section), an iterative strategy was implemented in which section z was aligned to section z+1. This approach preserves the continuity of deformation and minimizes global drift. The rigid registration was defined as a 2D rotation and translation. A custom objective function was formulated to compute the sum of squared distances between nearest-neighbor nuclei in adjacent slices. Nearest neighbors were identified using a KD-tree search, and the optimal transformation parameters were obtained by minimizing this objective using the minimize function from the SciPy optimization library in Python. Registration quality was visually inspected between adjacent sections, and minor manual adjustments were applied between slices z9 and z10 as well as z16 and z17 to improve local alignment. After registration, the entire 3D model was reoriented such that the anterior side faces left and the dorsal side faces upward. Finally, visualization of the 3D point cloud model was performed using the plotly package (v4.10.4) in R. Gene expression levels or annotated layers were projected onto the 3D model to enable spatial visualization. To reduce visual occlusion of internal points, particularly in the lateral view, a small upward jitter along the z axis was added to each point, and point opacity was set to 75%.

To generate smooth 3D surfaces (Fig. 5a) from spatial point cloud data, alpha shapes were computed for each anatomical region using the alphashape3d package (v1.3.2) in R. The resulting shapes were converted into triangular meshes and further smoothed using mesh processing functions from the Rvcg package (v0.24). Meshes were exported in PLY format, adjusted in Blender software (v4.3) to correct face orientation, and re-imported into R for visualization. Final rendering was performed using the rgl package (v1.3.17), with appropriate lighting and transparency settings to enhance structural clarity.

### 7. Cross-species and cross-modality analysis

Cross-species and cross-modality comparison was performed using SAMap^26^ (v1.0.15), which generated transcriptomic alignments between homologous cell groups across the two datasets. The resulting alignment scores were visualized as heatmaps using the ComplexHeatmap^72^ R package (v2.22.0). In this study, various combinations of cell groups, such as clusters or tissue-defined regions, from different datasets (e.g., amphioxus vs. mouse; snRNA-seq vs. Xenium) were selected for pairwise comparisons. The default SAMap pipeline relies on reciprocal BLAST between the gene sets of the two datasets, using a relatively permissive e-value threshold of 1 × 10^-6^ to identify putative homologs. However, this approach may lead to cell group alignments driven by gene pairs that are not truly homologous or functionally related. To mitigate the inclusion of spurious gene pairs, OrthoFinder^73^ (v2.5.4) was used to identify high-confidence orthologous relationships between two species. Only gene pairs supported by OrthoFinder were retained in the SAMap gene-pair correspondence tables for downstream alignment. In the amphioxus snRNA-seq versus Xenium comparison, the SAMap gene-pair correspondence tables were limited to one-to-one identity matches between genes with the same gene ID, given that both datasets originated from the same species.

## Acknowledgements

The authors thank Dr. Sean Tan, Kok Ting Chong, Sharon Gwee, Dr. Quy Xiao Xuan Lin, and Kok Chuan Tan from 10x Genomics, as well as Owen Kuo and Johnson Chen from Pharmigene, for their assistance with the Xenium experiments. We thank Ya-Ching Kuo and Tzu-Kai Huang from the Institute of Cellular and Organismic Biology (ICOB) for animal husbandry and maintenance, and Dr. Hsin-Ju Chuang from ICOB for help with paraffin embedding. We also thank Dr. Chun-Shiu Wu and Kai-Xuan Ding from the High-Throughput Genomics Core for assistance with sample quality control, and Shin-Yi Du and Dr. Wei-Chen Chu from ICOB for help with imaging. We are grateful to Marcus Calkins for English editing, and to Dr. Jason Leong and Dr. Tsai-Ming Lu for valuable discussions. Finally, we deeply appreciate the insightful comments and suggestions from Dr. Linda Z. Holland. This work was supported by grants 113-2621-B-001-004-MY3 from the National Science and Technology Council, Taiwan, and AS-GC-111-L01 from Academia Sinica, Taiwan.

## Author contributions

J.K.Y. supervised the project. J.K.Y. and C.Y.L. conceived and designed the study. C.Y.L., W.H.H., Y.H.C and Y.C.C. performed the experiments. C.Y.L. analyzed the data. J.K.Y. and C.Y.L. wrote the manuscript with input from Y.H.S., S.J.C., M.Y.L., and W.H.H.

## Reference

1 Sugahara, F., Murakami, Y., Pascual-Anaya, J. & Kuratani, S. Reconstructing the ancestral vertebrate brain. Dev Growth Differ 59, 163–174 (2017). 10.1111/dgd.12347

2 Kiecker, C. & Lumsden, A. The role of organizers in patterning the nervous system. Annu Rev Neurosci 35, 347–367 (2012). 10.1146/annurev-neuro-062111-150543

3 Sugahara, F. et al. Evidence from cyclostomes for complex regionalization of the ancestral vertebrate brain. Nature 531, 97–100 (2016). 10.1038/nature16518

4 Murakami, Y., Uchida, K., Rijli, F. M. & Kuratani, S. Evolution of the brain developmental plan: Insights from agnathans. Dev Biol 280, 249–259 (2005). 10.1016/j.ydbio.2005.02.008

5 Holland, L. Z. The origin and evolution of chordate nervous systems. Philos Trans R Soc Lond B Biol Sci 370 (2015). 10.1098/rstb.2015.0048

6 Satoh, N., Rokhsar, D. & Nishikawa, T. Chordate evolution and the three-phylum system. Proc Biol Sci 281, 20141729 (2014). 10.1098/rspb.2014.1729

7 Yong, L. W. et al. Somite Compartments in Amphioxus and Its Implications on the Evolution of the Vertebrate Skeletal Tissues. Front Cell Dev Biol 9, 607057 (2021). 10.3389/fcell.2021.607057

8 Putnam, N. H. et al. The amphioxus genome and the evolution of the chordate karyotype. Nature 453, 1064–1071 (2008). 10.1038/nature06967

9 Castro, A., Becerra, M., Manso, M. J. & Anadon, R. Neuronal Organization of the Brain in the Adult Amphioxus (Branchiostoma lanceolatum): A Study With Acetylated Tubulin Immunohistochemistry. J Comp Neurol 523, 2211–2232 (2015). 10.1002/cne.23785

10 Candiani, S., Moronti, L., Ramoino, P., Schubert, M. & Pestarino, M. A neurochemical map of the developing amphioxus nervous system. BMC Neurosci 13, 59 (2012). 10.1186/1471-2202-13-59

11 Wicht, H. & Lacalli, T. C. The nervous system of amphioxus: structure, development, and evolutionary significance. Canadian Journal of Zoology 83, 122–150 (2005). 10.1139/z04-163

12 Lacalli, T. C. Frontal eye circuitry, rostral sensory pathways and brain organization in amphioxus larvae: Evidence from 3D reconstructions. Philos T R Soc B 351, 243–263 (1996). DOI 10.1098/rstb.1996.0022

13 Lacalli, T. C., Holland, N. D. & West, J. E. Landmarks in the Anterior Central-Nervous-System of Amphioxus Larvae. Philos T R Soc B 344, 165–185 (1994). DOI 10.1098/rstb.1994.0059

14 Lacalli, T. C. & Kelly, S. J. Floor plate, glia and other support cells in the anterior nerve cord of amphioxus larvae. Acta Zool-Stockholm 83, 87–98 (2002). 10.1046/j.1463-6395.2002.00101.x

15 Vopalensky, P. et al. Molecular analysis of the amphioxus frontal eye unravels the evolutionary origin of the retina and pigment cells of the vertebrate eye. Proc Natl Acad Sci U S A 109, 15383–15388 (2012). 10.1073/pnas.1207580109

16 Pergner, J. & Kozmik, Z. Amphioxus photoreceptors - insights into the evolution of vertebrate opsins, vision and circadian rhythmicity. Int J Dev Biol 61, 665–681 (2017). 10.1387/ijdb.170230zk

17 Albuixech-Crespo, B. et al. Molecular regionalization of the developing amphioxus neural tube challenges major partitions of the vertebrate brain. Plos Biol 15, e2001573 (2017). 10.1371/journal.pbio.2001573

18. Candiani, S. & Pestarino, M. in Oxford Research Encyclopedia of Neuroscience (2018).

19 Holland, L. Z. Cephalochordata. Evolutionary Developmental Biology of Invertebrates 6, 91–133 (2015).

20 Benito-Gutierrez, E. et al. The dorsoanterior brain of adult amphioxus shares similarities in expression profile and neuronal composition with the vertebrate telencephalon. BMC Biol 19, 110 (2021). 10.1186/s12915-021-01045-w

21 Luecken, M. D. & Theis, F. J. Current best practices in single-cell RNA-seq analysis: a tutorial. Mol Syst Biol 15, e8746 (2019). 10.15252/msb.20188746

22 Lamanna, F. et al. A lamprey neural cell type atlas illuminates the origins of the vertebrate brain. Nat Ecol Evol 7, 1714–1728 (2023). 10.1038/s41559-023-02170-1

23 Castro, A., Becerra, M., Manso, M. J., Sherwood, N. M. & Anadon, R. Anatomy of the Hesse photoreceptor cell axonal system in the central nervous system of amphioxus. J Comp Neurol 494, 54–62 (2006). 10.1002/cne.20783

24 Ma, P. et al. Joint profiling of gene expression and chromatin accessibility during amphioxus development at single-cell resolution. Cell Rep 39, 110979 (2022). 10.1016/j.celrep.2022.110979

25 Ferran, J. L. & Puelles, L. Lessons from Amphioxus Bauplan About Origin of Cranial Nerves of Vertebrates That Innervates Extrinsic Eye Muscles. Anat Rec (Hoboken) 302, 452–462 (2019). 10.1002/ar.23824

26 Tarashansky, A. J. et al. Mapping single-cell atlases throughout Metazoa unravels cell type evolution. Elife 10 (2021). 10.7554/eLife.66747

27 Bozzo, M. et al. Amphioxus neuroglia: Molecular characterization and evidence for early compartmentalization of the developing nerve cord. Glia 69, 1654–1678 (2021). 10.1002/glia.23982

28 Lacalli, T. Amphioxus, motion detection, and the evolutionary origin of the vertebrate retinotectal map. Evodevo 9, 6 (2018). 10.1186/s13227-018-0093-2

29 Sterba, G., Fredriksson, G. & Olsson, R. Immunocytochemical Investigations of the Infundibular Organ in Amphioxus (Branchiostoma lanceolatum; Cephalochordata). Acta Zool-Stockholm 64, 149–153 (2010). 10.1111/j.1463-6395.1983.tb00793.x

30 Guerra, M. M. et al. Understanding How the Subcommissural Organ and Other Periventricular Secretory Structures Contribute via the Cerebrospinal Fluid to Neurogenesis. Front Cell Neurosci 9, 480 (2015). 10.3389/fncel.2015.00480

31 Castro, L. F., Rasmussen, S. L., Holland, P. W., Holland, N. D. & Holland, L. Z. A Gbx homeobox gene in amphioxus: insights into ancestry of the ANTP class and evolution of the midbrain/hindbrain boundary. Dev Biol 295, 40–51 (2006). 10.1016/j.ydbio.2006.03.003

32 La Manno, G. et al. Molecular architecture of the developing mouse brain. Nature 596, 92–96 (2021). 10.1038/s41586-021-03775-x

33 Shi, Y. et al. Decoding the spatiotemporal regulation of transcription factors during human spinal cord development. Cell Res 34, 193–213 (2024). 10.1038/s41422-023-00897-x

34 Wilson, L. & Maden, M. The mechanisms of dorsoventral patterning in the vertebrate neural tube. Dev Biol 282, 1–13 (2005). 10.1016/j.ydbio.2005.02.027

35 Shimamura, K., Hartigan, D. J., Martinez, S., Puelles, L. & Rubenstein, J. L. Longitudinal organization of the anterior neural plate and neural tube. Development 121, 3923–3933 (1995). 10.1242/dev.121.12.3923

36 Pani, A. M. et al. Ancient deuterostome origins of vertebrate brain signalling centres. Nature 483, 289–294 (2012). 10.1038/nature10838

37 Subramanian, L. & Tole, S. Mechanisms underlying the specification, positional regulation, and function of the cortical hem. Cereb Cortex 19 Suppl 1, i90–95 (2009). 10.1093/cercor/bhp031

38 Caronia-Brown, G., Yoshida, M., Gulden, F., Assimacopoulos, S. & Grove, E. A. The cortical hem regulates the size and patterning of neocortex. Development 141, 2855–2865 (2014). 10.1242/dev.106914

39 Subramanian, L., Remedios, R., Shetty, A. & Tole, S. Signals from the edges: the cortical hem and antihem in telencephalic development. Semin Cell Dev Biol 20, 712–718 (2009). 10.1016/j.semcdb.2009.04.001

40 Kiecker, C. & Lumsden, A. Hedgehog signaling from the ZLI regulates diencephalic regional identity. Nat Neurosci 7, 1242–1249 (2004). 10.1038/nn1338

41 Holland, L. Z. et al. Evolution of bilaterian central nervous systems: a single origin? Evodevo 4 (2013). Artn 27 10.1186/2041-9139-4-27

42 Holland, L. Z. & Holland, N. D. Cephalochordates: A window into vertebrate origins. Curr Top Dev Biol 141, 119–147 (2021). 10.1016/bs.ctdb.2020.07.001

43 Toresson, H., Martinez-Barbera, J. P., Bardsley, A., Caubit, X. & Krauss, S. Conservation of BF-1 expression in amphioxus and zebrafish suggests evolutionary ancestry of anterior cell types that contribute to the vertebrate telencephalon. Dev Genes Evol 208, 431–439 (1998). 10.1007/s004270050200

44 Puelles, L. et al. Pallial and subpallial derivatives in the embryonic chick and mouse telencephalon, traced by the expression of the genes Dlx-2, Emx-1, Nkx-2.1, Pax-6, and Tbr-1. J Comp Neurol 424, 409–438 (2000). 10.1002/1096-9861(20000828)424:3<409::aid-cne3>3.0.co;2-7

45 Dai, Y. et al. Evolutionary origin of the chordate nervous system revealed by amphioxus developmental trajectories. Nat Ecol Evol 8, 1693–1710 (2024). 10.1038/s41559-024-02469-7

46 Glenn Northcutt, R. The new head hypothesis revisited. J Exp Zool B Mol Dev Evol 304, 274–297 (2005). 10.1002/jez.b.21063

47 Zhang, C. L., Zou, Y., Yu, R. T., Gage, F. H. & Evans, R. M. Nuclear receptor TLX prevents retinal dystrophy and recruits the corepressor atrophin1. Genes Dev 20, 1308–1320 (2006). 10.1101/gad.1413606

48 Coomer, C. E. et al. Her9/Hes4 is required for retinal photoreceptor development, maintenance, and survival. Sci Rep 10, 11316 (2020). 10.1038/s41598-020-68172-2

49 Yip, H. K. Retinal stem cells and regeneration of vision system. Anat Rec (Hoboken) 297, 137–160 (2014). 10.1002/ar.22800

50 Martinez-De Luna, R. I., Kelly, L. E. & El-Hodiri, H. M. The Retinal Homeobox (Rx) gene is necessary for retinal regeneration. Dev Biol 353, 10–18 (2011). 10.1016/j.ydbio.2011.02.008

51 Lord, M. N., Subramanian, K., Kanoski, S. E. & Noble, E. E. Melanin-concentrating hormone and food intake control: Sites of action, peptide interactions, and appetition. Peptides 137, 170476 (2021). 10.1016/j.peptides.2020.170476

52 Yamaguchi, Y. Arginine vasopressin: Critical regulator of circadian homeostasis. Peptides 177, 171229 (2024). 10.1016/j.peptides.2024.171229

53 Gomez, M. D., Angueyra, J. M. & Nasi, E. Light-transduction in melanopsin-expressing photoreceptors of Amphioxus. P Natl Acad Sci USA 106, 9081–9086 (2009). 10.1073/pnas.0900708106

54 Koyanagi, M., Kubokawa, K., Tsukamoto, H., Shichida, Y. & Terakita, A. Cephalochordate melanopsin: evolutionary linkage between invertebrate visual cells and vertebrate photosensitive retinal ganglion cells. Curr Biol 15, 1065–1069 (2005). 10.1016/j.cub.2005.04.063

55 Ferrer, C., Malagon, G., Gomez Mdel, P. & Nasi, E. Dissecting the determinants of light sensitivity in amphioxus microvillar photoreceptors: possible evolutionary implications for melanopsin signaling. J Neurosci 32, 17977–17987 (2012). 10.1523/JNEUROSCI.3069-12.2012

56 Ekhart, D., Korf, H. W. & Wicht, H. Cytoarchitecture, topography, and descending supraspinal projections in the anterior central nervous system of Branchiostoma lanceolatum. J Comp Neurol 466, 319–330 (2003). 10.1002/cne.10803

57 Nilaweera, K. N., Ellis, C., Barrett, P., Mercer, J. G. & Morgan, P. J. Hypothalamic bHLH transcription factors are novel candidates in the regulation of energy balance. Eur J Neurosci 15, 644–650 (2002). 10.1046/j.1460-9568.2002.01894.x

58 Mazet, F. & Shimeld, S. M. The evolution of chordate neural segmentation. Dev Biol 251, 258–270 (2002). 10.1006/dbio.2002.0831

59 Benito-Gutierrez, E. A gene catalogue of the amphioxus nervous system. Int J Biol Sci 2, 149–160 (2006). 10.7150/ijbs.2.149

60 Simakov, O. et al. Deeply conserved synteny resolves early events in vertebrate evolution. Nat Ecol Evol 4, 820–830 (2020). 10.1038/s41559-020-1156-z

61 Gattoni, G., Keitley, D., Sawle, A. & Benito-Gutierrez, E. An ancient apical patterning system sets the position of the forebrain in chordates. Sci Adv 11, eadq4731 (2025). 10.1126/sciadv.adq4731

62 Fleming, S. J. et al. Unsupervised removal of systematic background noise from droplet-based single-cell experiments using CellBender. Nat Methods 20, 1323–1335 (2023). 10.1038/s41592-023-01943-7

63 Hao, Y. et al. Dictionary learning for integrative, multimodal and scalable single-cell analysis. Nat Biotechnol (2023). 10.1038/s41587-023-01767-y

64 Stuart, T. et al. Comprehensive Integration of Single-Cell Data. Cell 177, 1888–1902 e1821 (2019). 10.1016/j.cell.2019.05.031

65 Germain, P. L., Lun, A., Garcia Meixide, C., Macnair, W. & Robinson, M. D. Doublet identification in single-cell sequencing data using scDblFinder. F1000Res 10, 979 (2021). 10.12688/f1000research.73600.2

66 Hua, Y., Weng, L., Zhao, F. & Rambow, F. SeuratExtend: streamlining single-cell RNA-seq analysis through an integrated and intuitive framework. Gigascience 14 (2025). 10.1093/gigascience/giaf076

67 Wolf, F. A., Angerer, P. & Theis, F. J. SCANPY: large-scale single-cell gene expression data analysis. Genome Biol 19, 15 (2018). 10.1186/s13059-017-1382-0

68 Conesa, A. & Gotz, S. Blast2GO: A comprehensive suite for functional analysis in plant genomics. Int J Plant Genomics 2008, 619832 (2008). 10.1155/2008/619832

69 Cantalapiedra, C. P., Hernandez-Plaza, A., Letunic, I., Bork, P. & Huerta-Cepas, J. eggNOG-mapper v2: Functional Annotation, Orthology Assignments, and Domain Prediction at the Metagenomic Scale. Mol Biol Evol 38, 5825–5829 (2021). 10.1093/molbev/msab293

70 Huerta-Cepas, J. et al. eggNOG 5.0: a hierarchical, functionally and phylogenetically annotated orthology resource based on 5090 organisms and 2502 viruses. Nucleic Acids Res 47, D309–D314 (2019). 10.1093/nar/gky1085

71 Wu, T. et al. clusterProfiler 4.0: A universal enrichment tool for interpreting omics data. Innovation (Camb) 2, 100141 (2021). 10.1016/j.xinn.2021.100141

72 Gu, Z., Eils, R. & Schlesner, M. Complex heatmaps reveal patterns and correlations in multidimensional genomic data. Bioinformatics 32, 2847–2849 (2016). 10.1093/bioinformatics/btw313

73 Emms, D. M. & Kelly, S. OrthoFinder: phylogenetic orthology inference for comparative genomics. Genome Biol 20, 238 (2019). 10.1186/s13059-019-1832-y

