## Supplementary Figures for "A 3D Amphioxus Brain Atlas Illuminates the Blueprint of the Ancestral Chordate Brain"

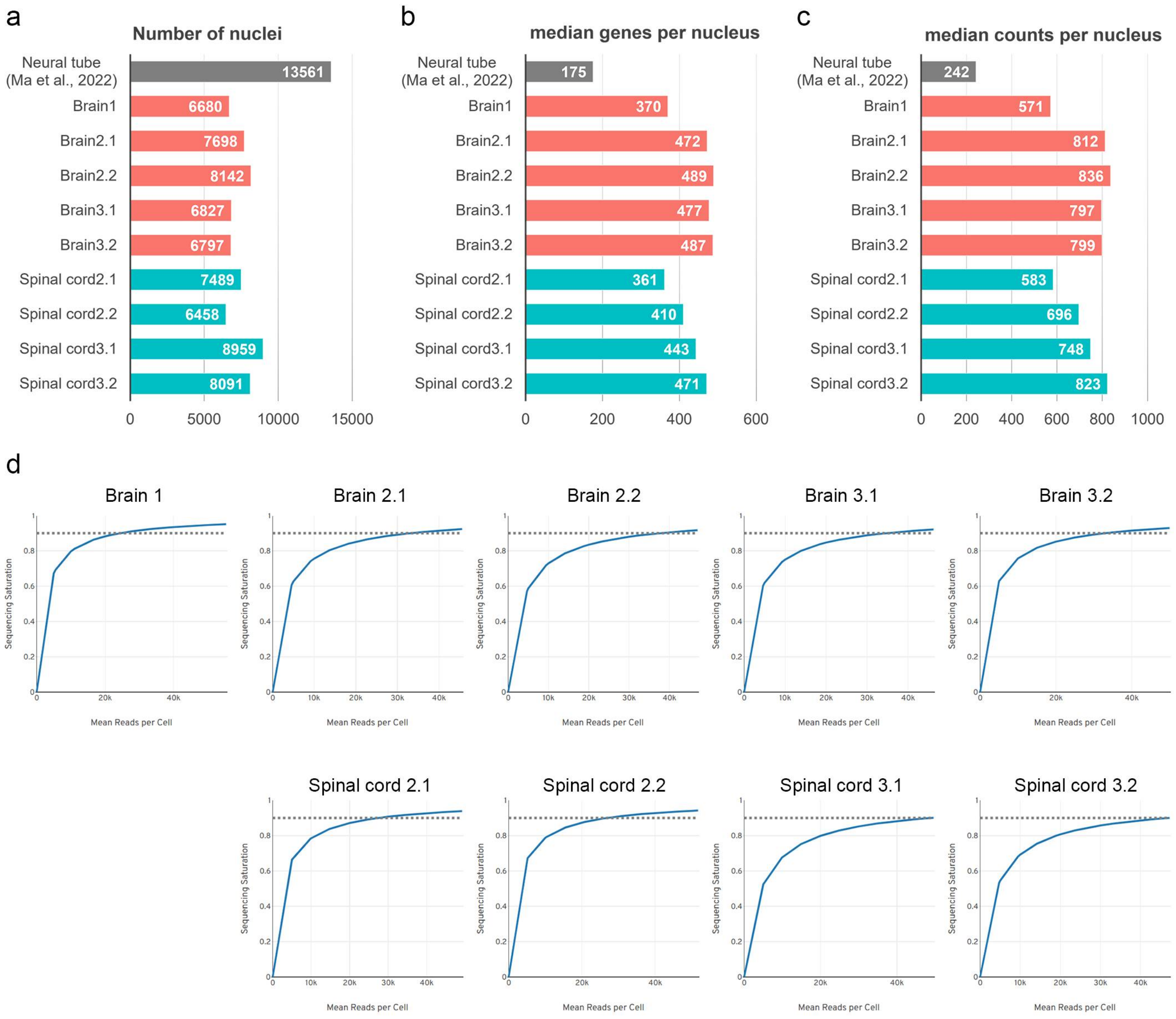


**Figure S1 | Statistics of snRNA sequencing data.**

**a-c**, Bar plots show the number of nuclei (**a**), median number of genes per nucleus (**b**), and median number of UMIs per nucleus (**c**). In the brain/spinal cord X.Y notation, X denotes biological replicates (independent batches) and Y denotes technical replicates. The metrics demonstrate improved sensitivity and overall data quality of our snRNA-seq dataset compared with a previous study[24]. **d**, Sequencing saturation curves obtained by downsampling the sequencing depth of each sample. The dotted line marks the approximate saturation point.

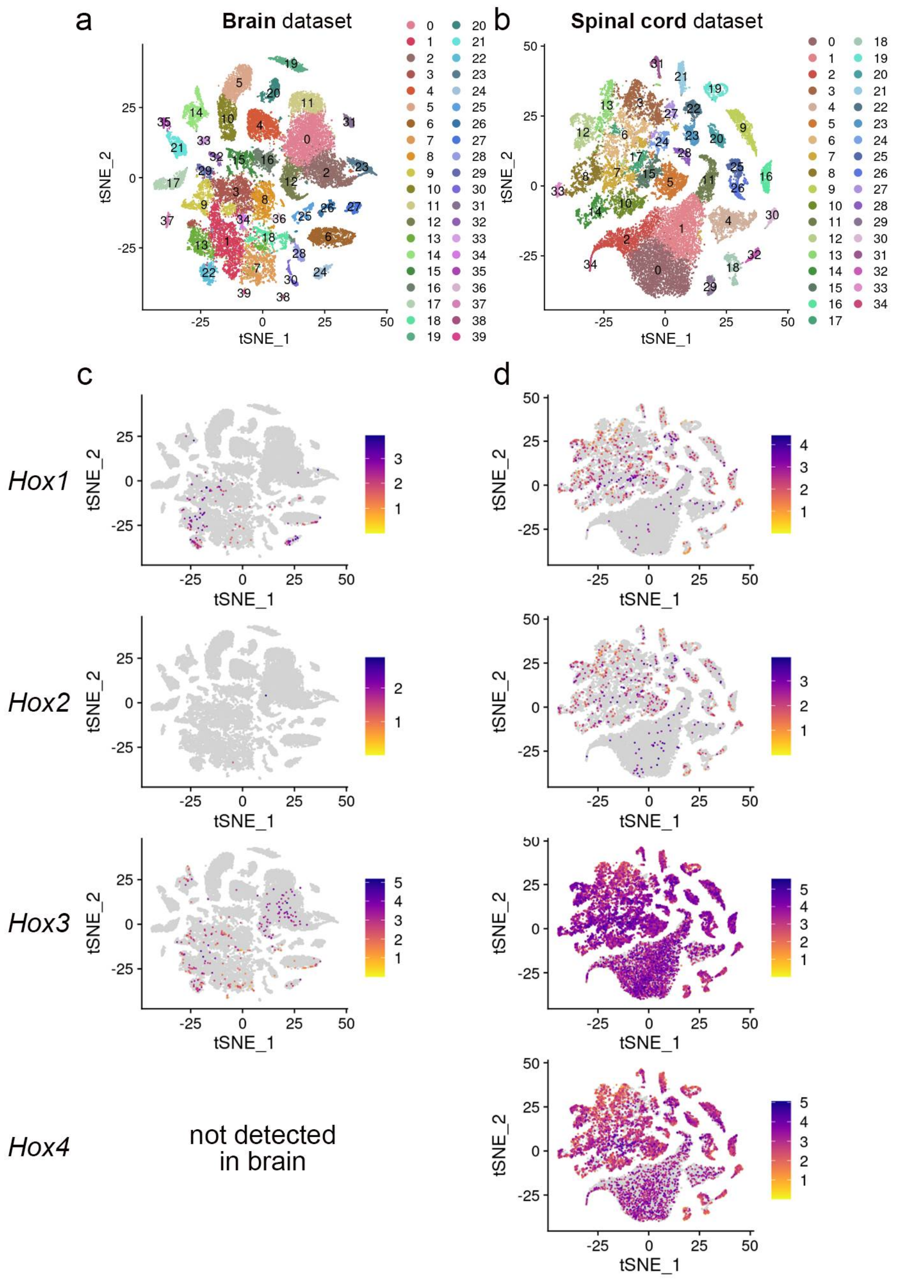


**Figure S2 | Expression profiles of *Hox* genes in brain and spinal cord snRNA-seq datasets.** **a-b**, t-SNE plots of nuclei from the brain (**a**) and spinal cord (**b**) with cluster labels. **c-d**, Feature plots of representative *Hox* genes showing minimal expression in the brain (**c**) but robust expression in the spinal cord (**d**). Results are consistent with the characteristic posterior spatial distribution of *Hox* gene expression during development[25].

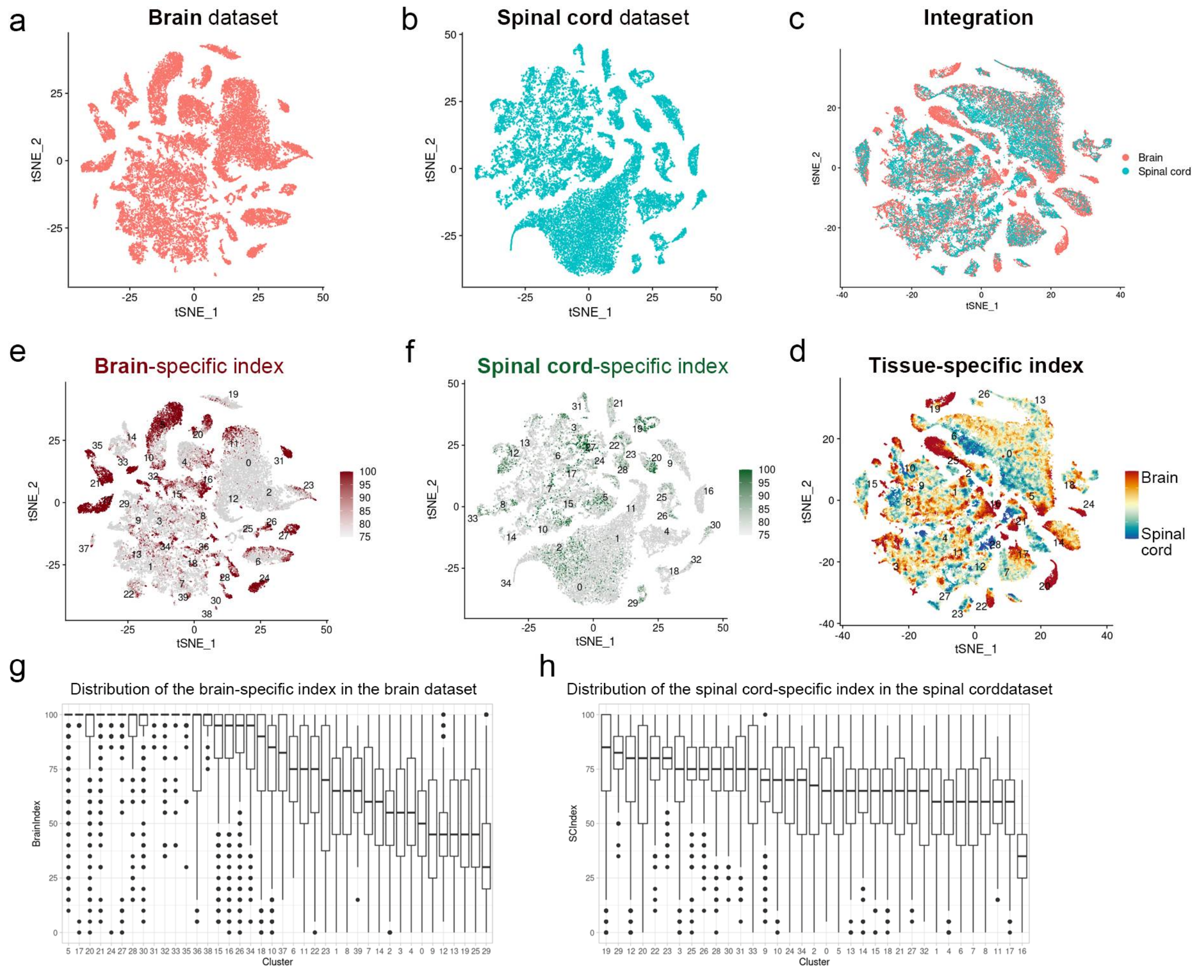


**Figure S3 | Tissue-specific index reveals molecular complexity in the brain snRNA-seq dataset.** **a-c**, t-SNE plots of nuclei from the brain (**a**) and spinal cord (**b**) and integrated (**c**) dataset. **d**, t-SNE plot of the integrated dataset colored by tissue-specific index. Red indicates brain-enriched, and blue indicates spinal cord-enriched profiles. **e-f**, Brain- and spinal cord-specific indices projected separately onto the separate t-SNE plots of brain (**e**) and spinal cord (**f**) datasets. **g-h**, Distribution of brain- and spinal cord-specific indices across clusters in the brain (**g**) and spinal cord (**h**) datasets, respectively.

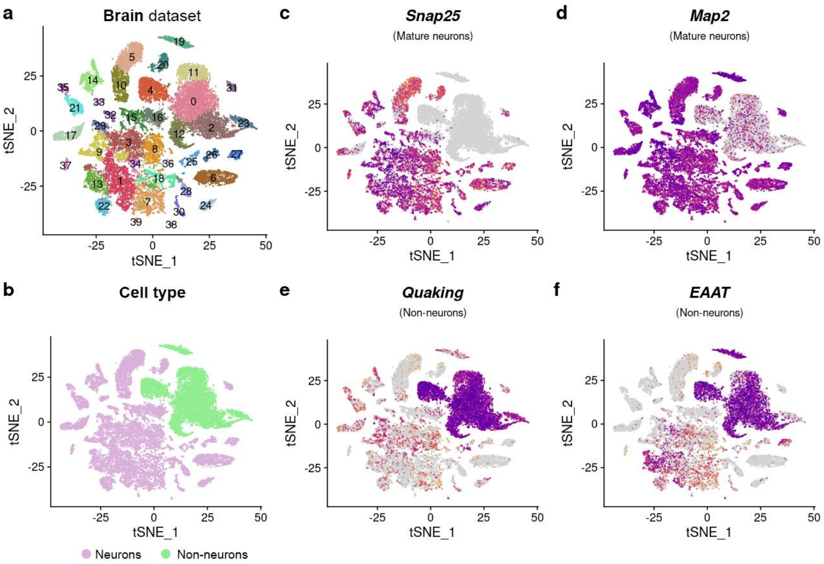


**Figure S4 | Expression profiles of neuronal and non-neuronal markers in the brain snRNA-seq dataset.**

**a**, t-SNE plot of the brain dataset with cluster labels. **b**, t-SNE plot colored by neuronal and non-neuronal cell-type identities. **c-f**, Feature plots showing expression of representative neuronal markers *Snap25* (**c**) and *Map2* (**d**), and non-neuronal markers *Quaking* (**e**) and *EAAT* (**f**).

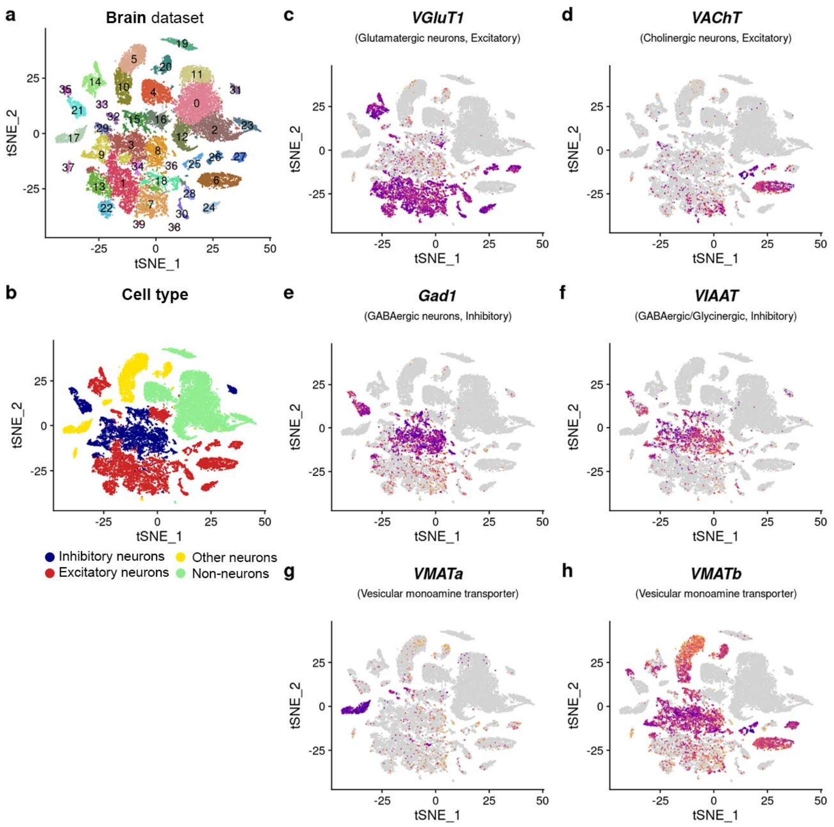


**Figure S5 | Expression profiles of neurotransmitter markers in the brain snRNA-seq dataset.** **a**, t-SNE plot of the brain dataset with cluster labels. **b**, t-SNE plot colored by major neurotransmitter classes. **c-h**, Feature plots showing the expression of excitatory neuron markers *VGluT1* (**c**) and *VAChT* (**d**), inhibitory neuron markers *Gad1* (**e**) and *VIAAT* (**f**), and monoaminergic transporters *VMATa* (**g**) and *VMATb* (**h**).

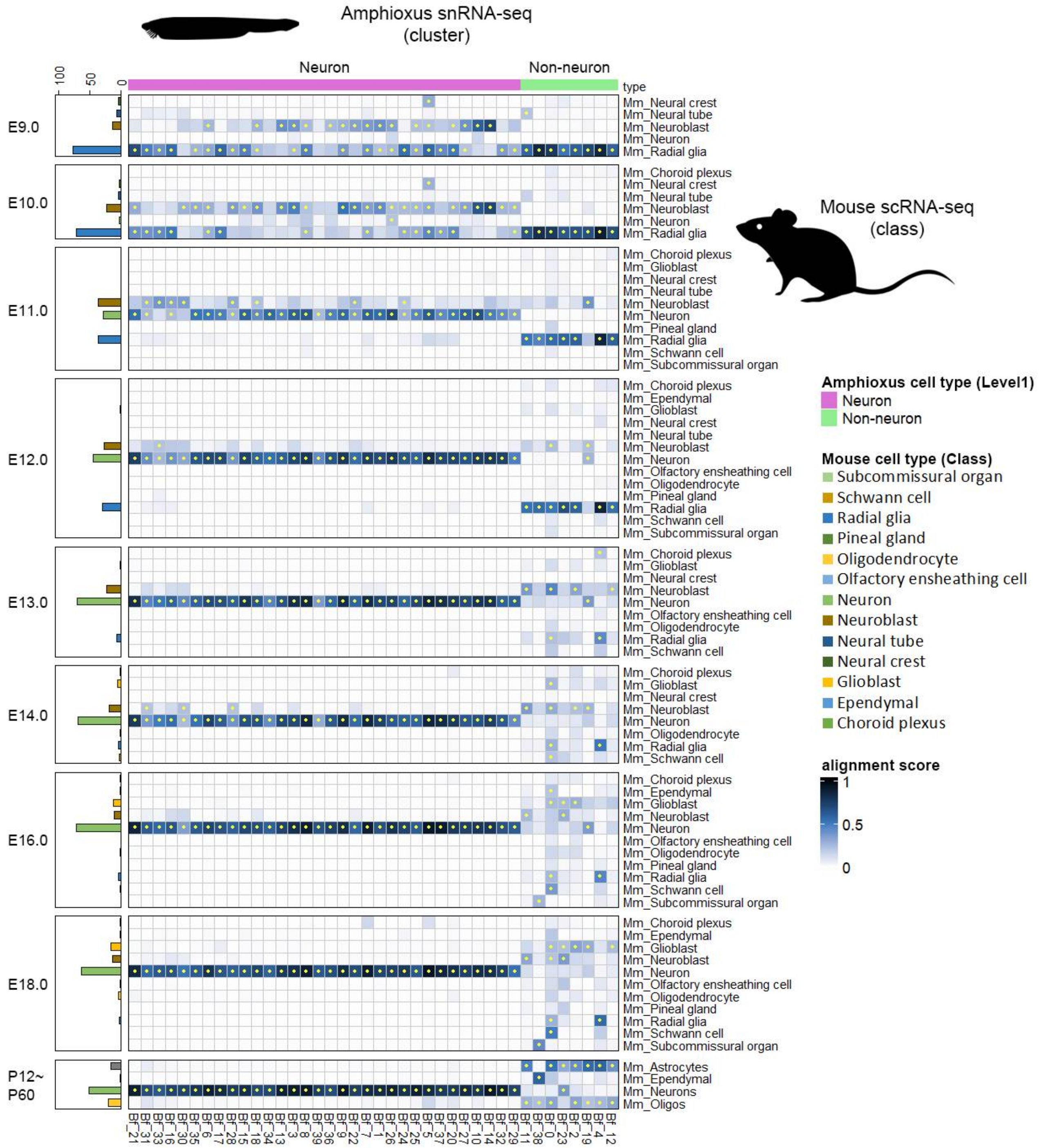


**Figure S6 | SAMap-based alignment of cell types between amphioxus and mouse brain datasets.** Heatmaps showing alignment scores derived from SAMap-based cross-species transcriptomic comparison. Each mouse dataset represents a distinct developmental stage (E9.0 to adolescence) and was independently aligned with the amphioxus dataset. Columns represent clusters from the amphioxus brain, and rows correspond to class-level annotations of the mouse brain defined in the original publication[32]. Non-neural mouse tissues, such as endoderm, mesoderm and blood, were excluded from the analysis. Higher alignment scores indicate higher transcriptomic similarity across species. Alignments with scores > 0.2 are marked with asterisks in the heatmap cells. Amphioxus neuronal (pink) and non-neuronal (green) clusters are indicated above the heatmap. Proportional bar charts on the left depict the relative abundance of different mouse cell types.

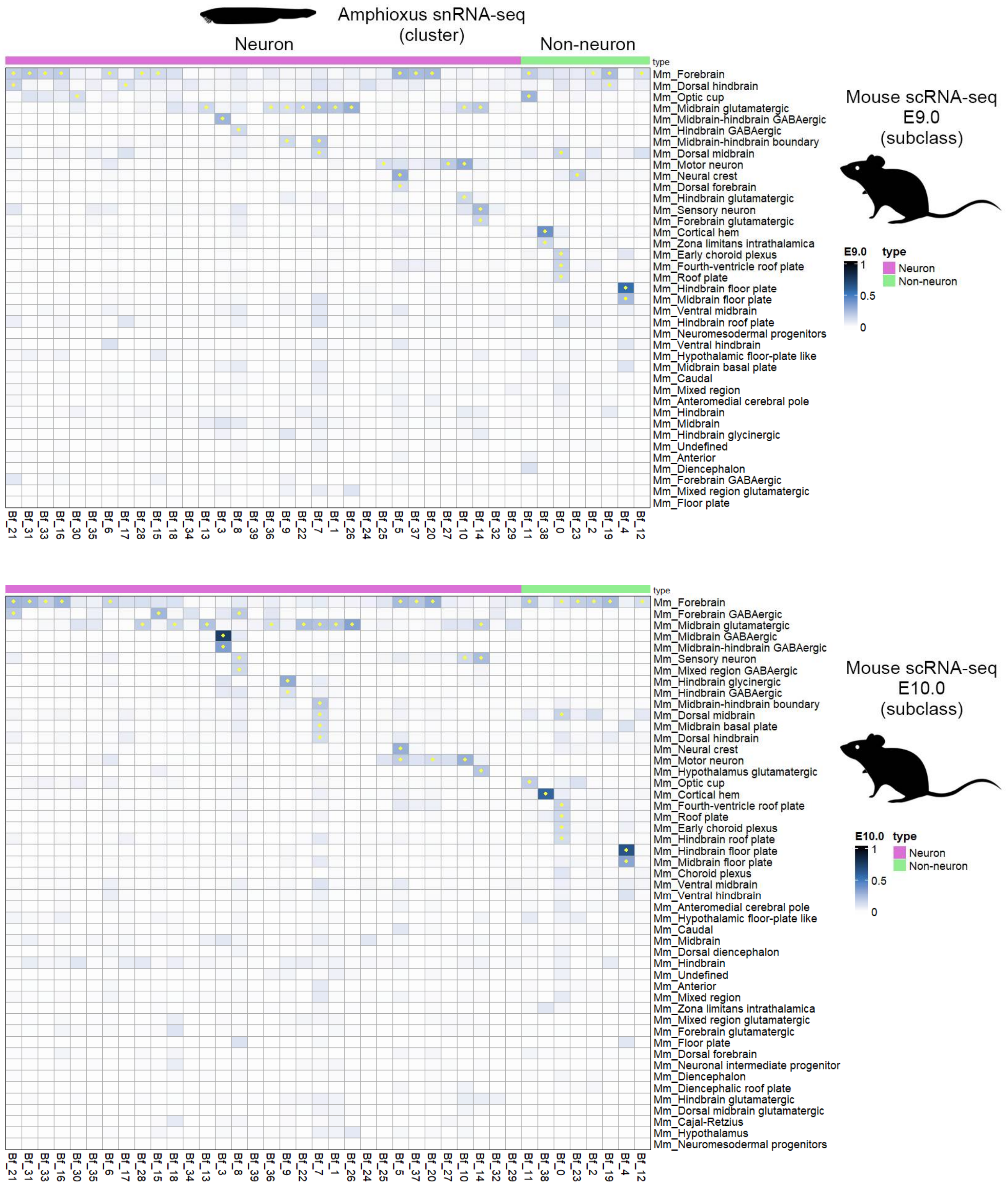


**Figure S7 | SAMap-based alignment of cell types between amphioxus and subclass-level annotations of E9.0 and E10.0 mouse brain datasets.**

Heatmaps showing alignment scores derived from SAMap-based cross-species transcriptomic comparison. Each mouse dataset represents a distinct developmental stage (E9.0 and E10.0) and was independently aligned with the amphioxus brain dataset. Columns represent clusters from the amphioxus brain, and rows correspond to subclass-level annotations of the mouse brain defined in the original publication[32]. Alignments with scores > 0.1 are marked with asterisks in the heatmap cells.

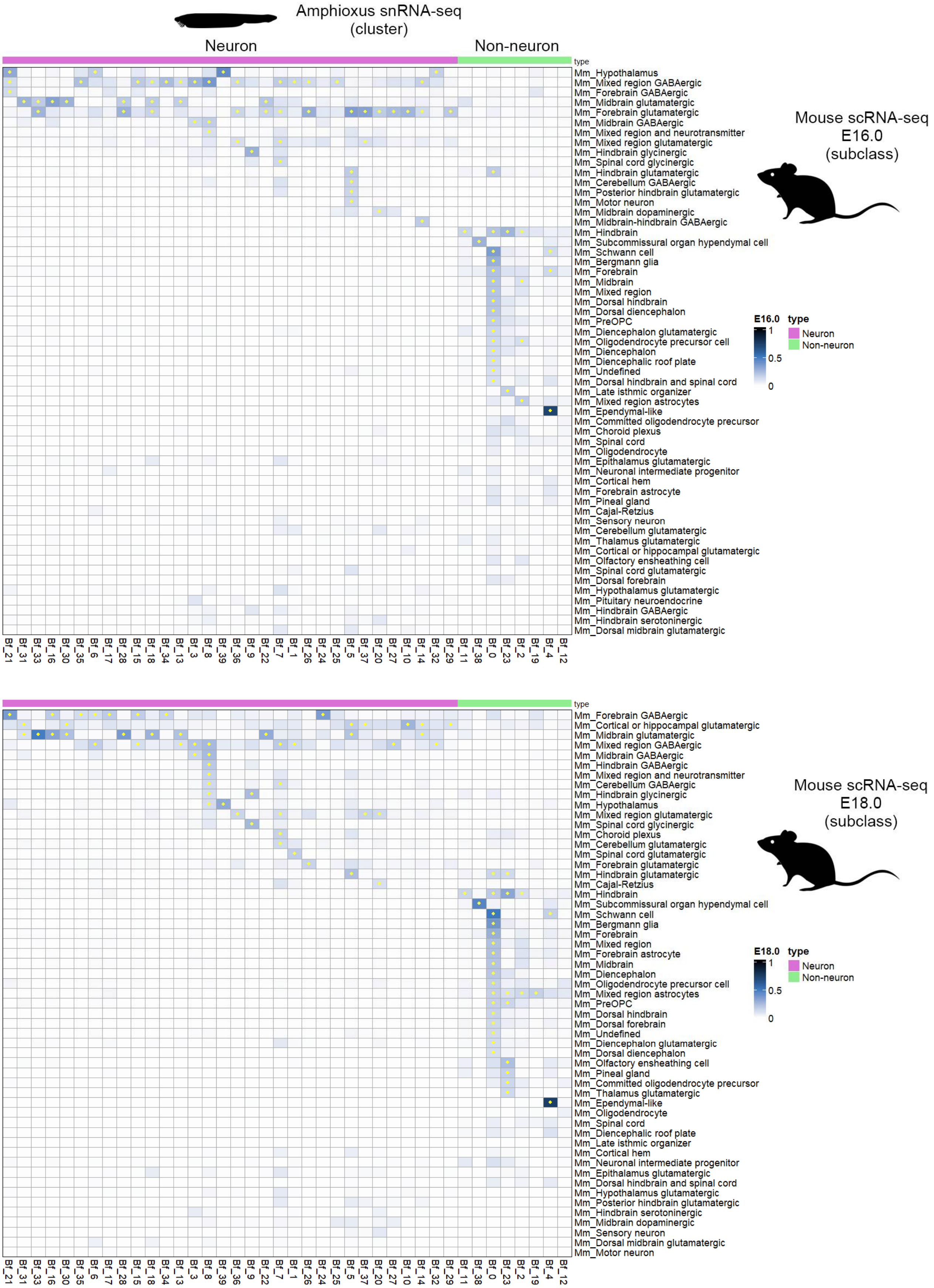


**Figure S8 | SAMap-based alignment of cell types between amphioxus and subclass-level annotations of E16.0 and E18.0 mouse brain datasets.**

Heatmaps showing alignment scores derived from SAMap-based cross-species transcriptomic comparison. Each mouse dataset represents a distinct developmental stage (E16.0 and E18.0) and was independently aligned with the amphioxus brain dataset. Columns represent clusters from the amphioxus brain, and rows correspond to subclass-level annotations of the mouse brain defined in the original publication[32]. Alignments with scores > 0.1 are marked with asterisks in the heatmap cells.

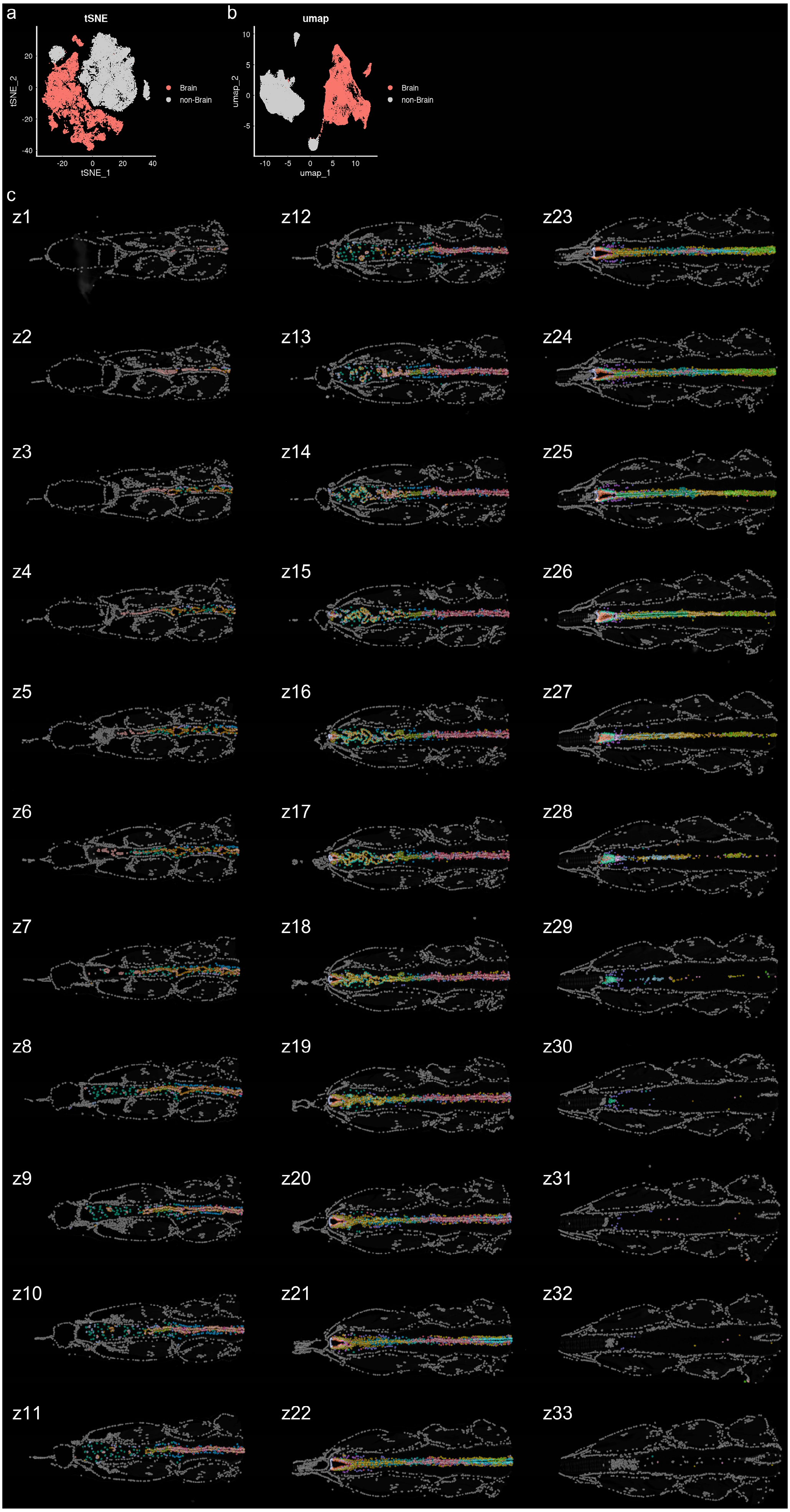


**Figure S9 | Full Xenium dataset with cell cluster information.**

**a-b**, t-SNE (a) and UMAP (b) plots of the Xenium dataset with brain cells in red and non-brain cells in grey. **c**, Spatial map of the full Xenium dataset across serial dorsal-ventral sections, oriented with anterior to the left, corresponding to Fig. 1f. Cells are colored according to their assigned clusters.

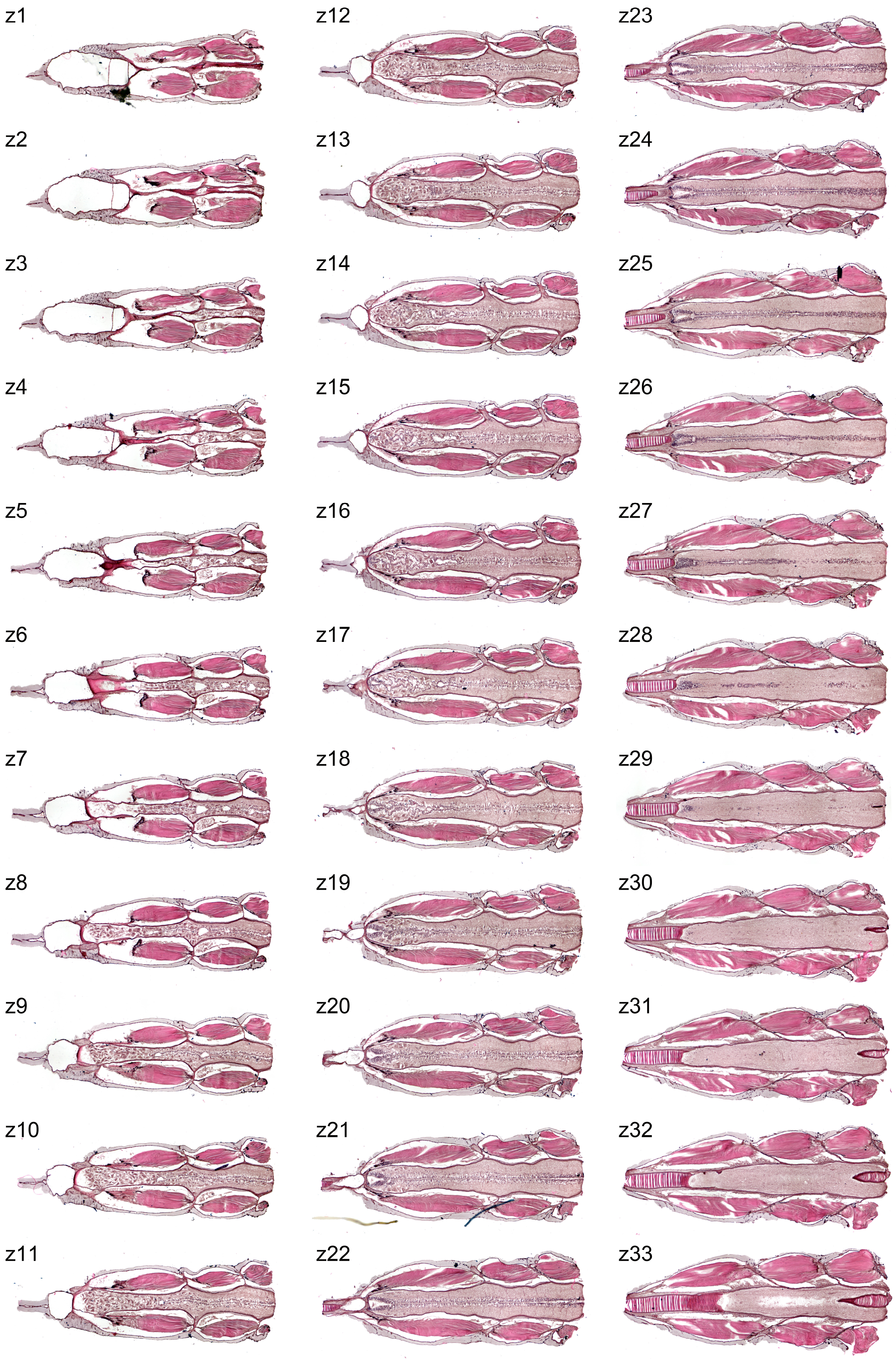


**Figure S10 | Full dataset of H&E staining of the Xenium slide.**

Post-H&E-stained tissues of the serial dorsal-ventral sections, oriented with anterior to the left. Data correspond to Fig. 1h.

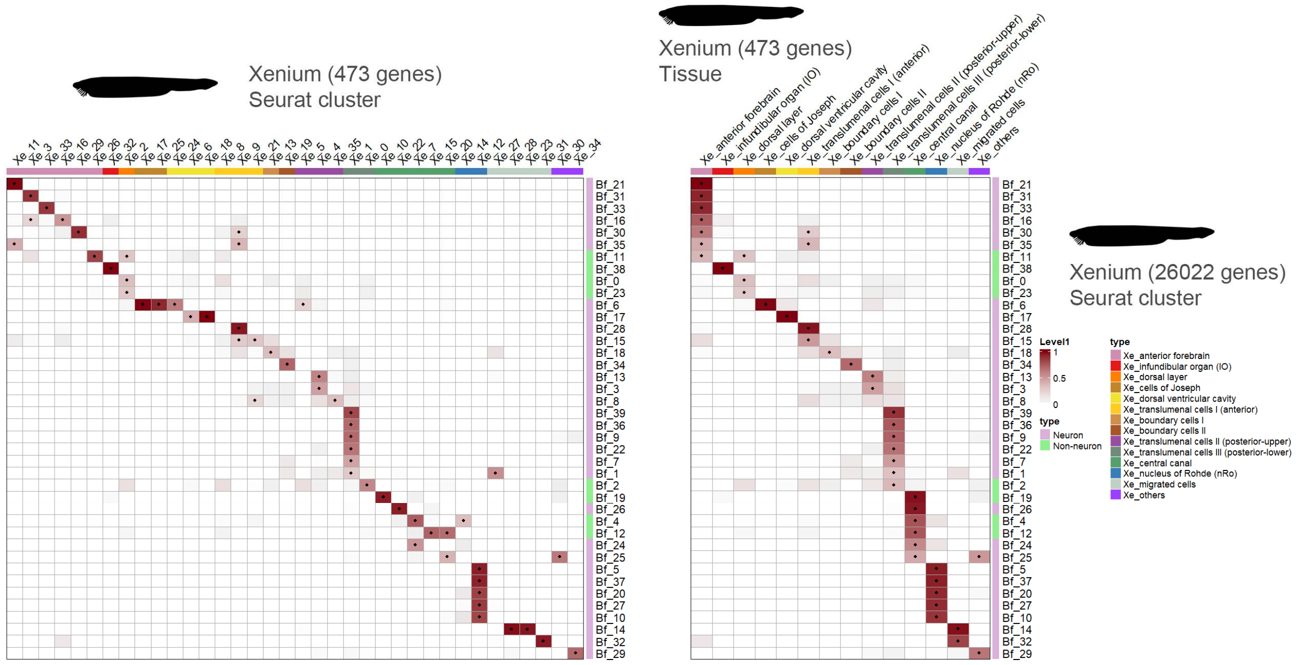


**Figure S11 | SAMap-based alignment of cell types between the amphioxus Xenium and snRNA-seq datasets.**

Heatmaps showing alignment scores derived from SAMap-based cross-modality transcriptomic comparison. The Xenium dataset analyses at the cluster level (left) and tissue level (right) were independently aligned with the snRNA-seq dataset. Columns represent clusters or tissues from the Xenium dataset, and rows correspond to cell clusters of the snRNA-seq dataset. Alignments with scores > 0.2 are marked with asterisks in the heatmap cells.

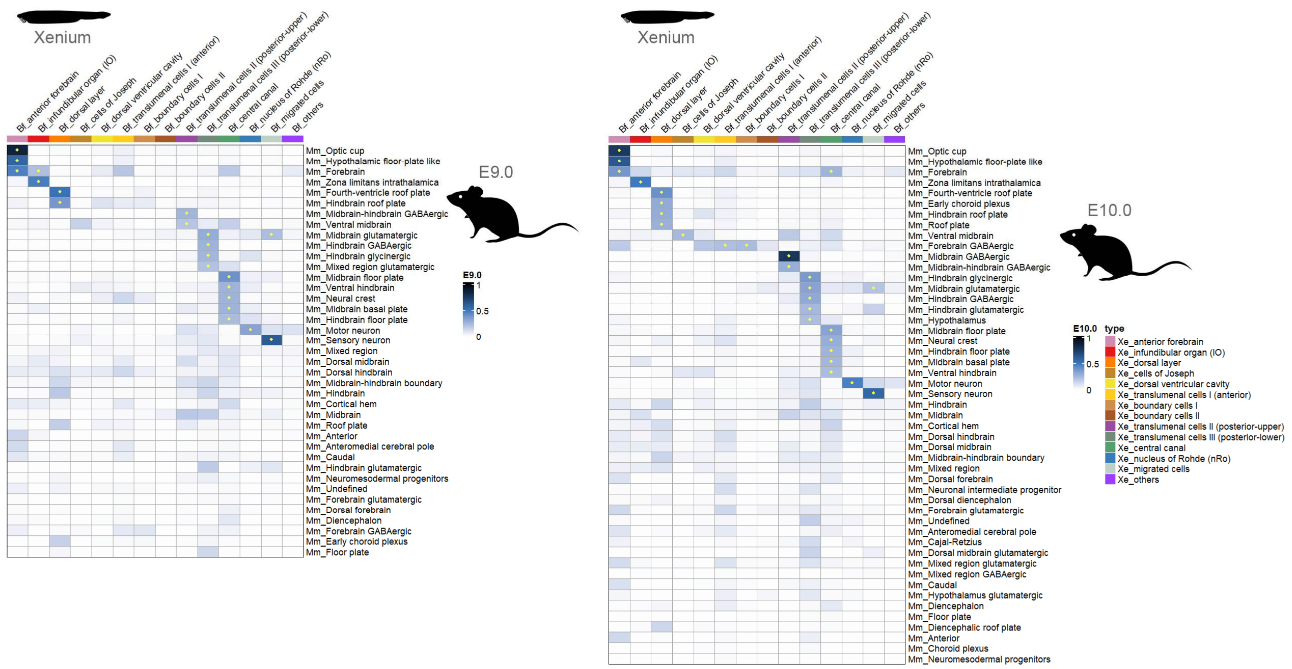


**Figure S12 | SAMap alignment scores between the amphioxus Xenium dataset and the E9.0 and E10.0 mouse brain datasets.**

Heatmaps showing SAMap-based cell-type alignments between the amphioxus Xenium dataset and mouse brain datasets at developmental stages E9.0 (left) and E10.0 (right)[32]. Columns represent tissue-level annotations from the amphioxus Xenium dataset, and rows correspond to subclass-level annotations of the mouse brain. Alignments with scores > 0.2 are marked with asterisks in the heatmap cells.

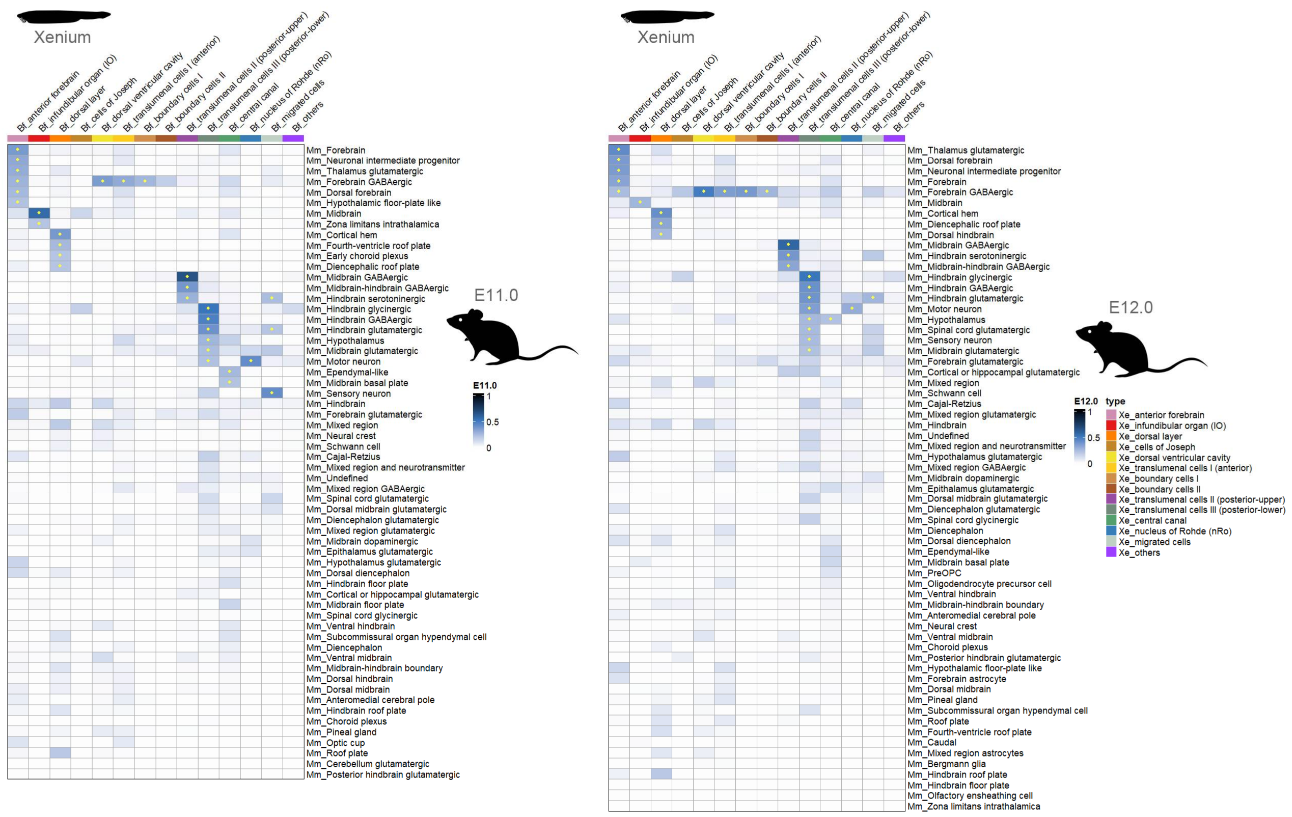


**Figure S13 | SAMap alignment scores between the amphioxus Xenium dataset and the E11.0 and E12.0 mouse brain datasets.**

Heatmaps showing SAMap-based cell-type alignments between the amphioxus Xenium dataset and mouse brain datasets at developmental stages E11.0 (left) and E12.0 (right)[32]. Columns represent tissue-level annotations from the amphioxus Xenium dataset, and rows correspond to subclass-level annotations of the mouse brain. Alignments with scores > 0.2 are marked with asterisks in the heatmap cells.

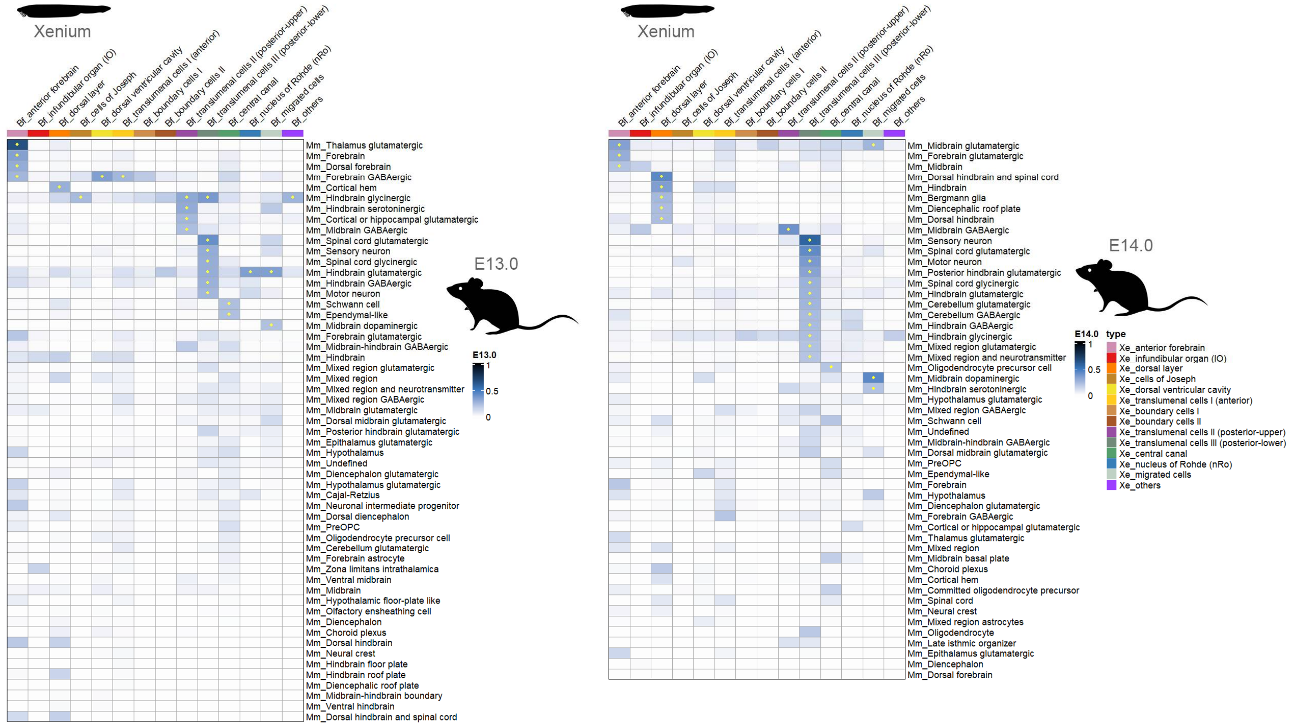


**Figure S14 | SAMap alignment scores between the amphioxus Xenium dataset and the E13.0 and E14.0 mouse brain datasets.**

Heatmaps showing SAMap-based cell-type alignments between the amphioxus Xenium dataset and mouse brain datasets at developmental stages E13.0 (left) and E14.0 (right)[32]. Columns represent tissue-level annotations from the amphioxus Xenium dataset, and rows correspond to subclass-level annotations of the mouse brain. Alignments with scores > 0.2 are marked with asterisks in the heatmap cells.

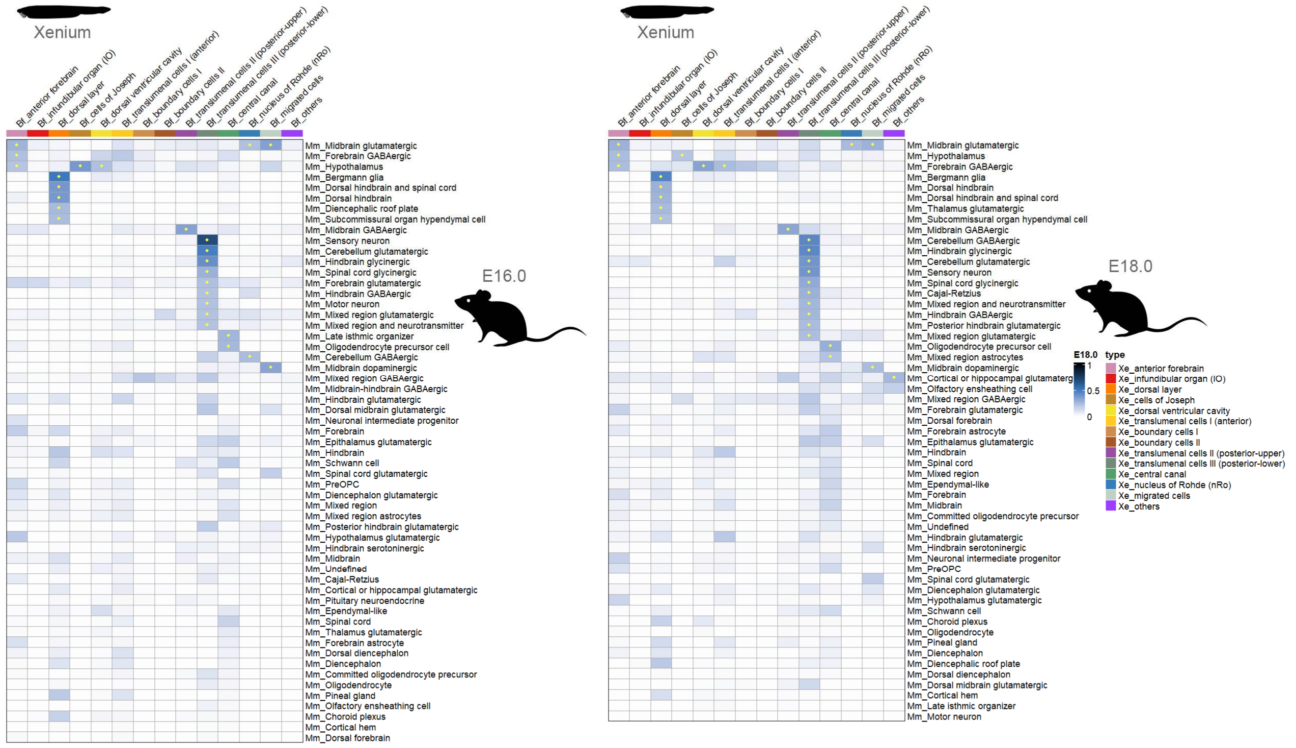


**Figure S15 | SAMap alignment scores between the amphioxus Xenium dataset and the E16.0 and E18.0 mouse brain datasets.**

Heatmaps showing SAMap-based cell-type alignments between the amphioxus Xenium dataset and mouse brain datasets at developmental stages E16.0 (left) and E18.0 (right)[32]. Columns represent tissue-level annotations from the amphioxus Xenium dataset, and rows correspond to subclass-level annotations of the mouse brain. Alignments with scores > 0.2 are marked with asterisks in the heatmap cells.

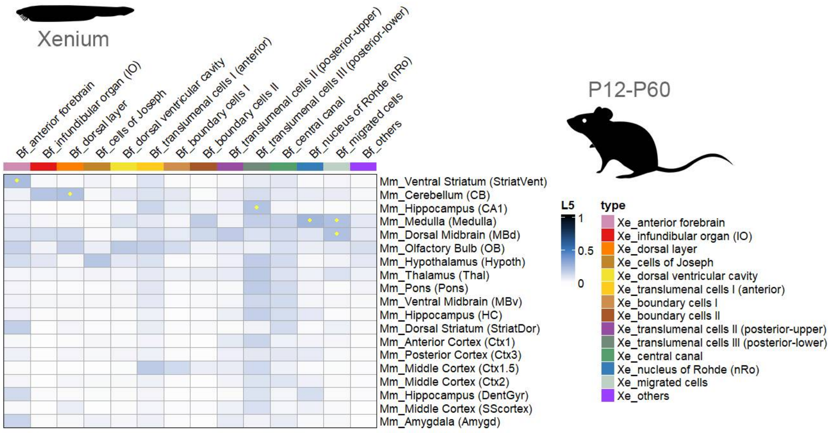


**Figure S16 | SAMap alignment scores between amphioxus the Xenium dataset and the adolescent mouse brain dataset.**

Heatmaps showing SAMap-based cell-type alignments between the amphioxus Xenium dataset and mouse brain dataset at adolescent stages[32]. Columns represent tissue-level annotations from the amphioxus Xenium dataset, and rows correspond to subclass-level annotations of the mouse brain. Alignments with scores > 0.2 are marked with asterisks in the heatmap cells.

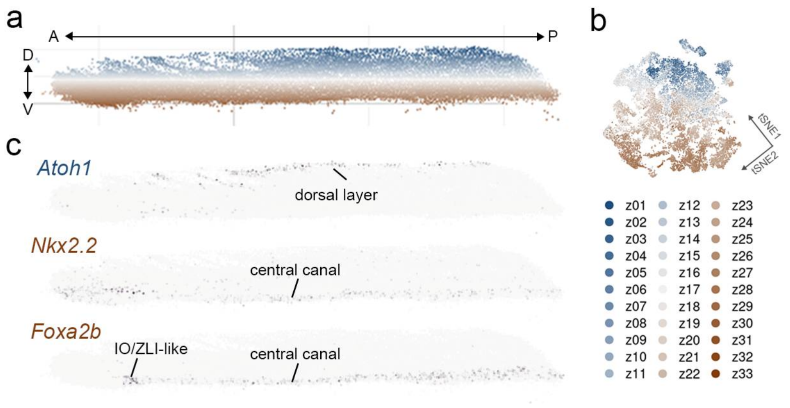


**Figure S17 | Dorsal-ventral (D-V) patterning of the adult amphioxus brain.**
**a**, Lateral view of the 3D point cloud model of the amphioxus brain, oriented with anterior to the left. Points are colored by section layers; dorsal sections are in sky blue and ventral sections are in earthy brown. **b**, Projection of section layers onto the *t*-SNE plot of the Xenium spatial transcriptomic dataset. **c**, Lateral view of the 3D point cloud model showing expression levels and domains of individual genes, including the roof plate marker *Atoh1* and floor plate markers *Nkx2.2* and *Foxa2b*. Color scales were normalized per gene, with the darkest color representing the maximal expression. Brain regions corresponding to gene expression domains were indicated.

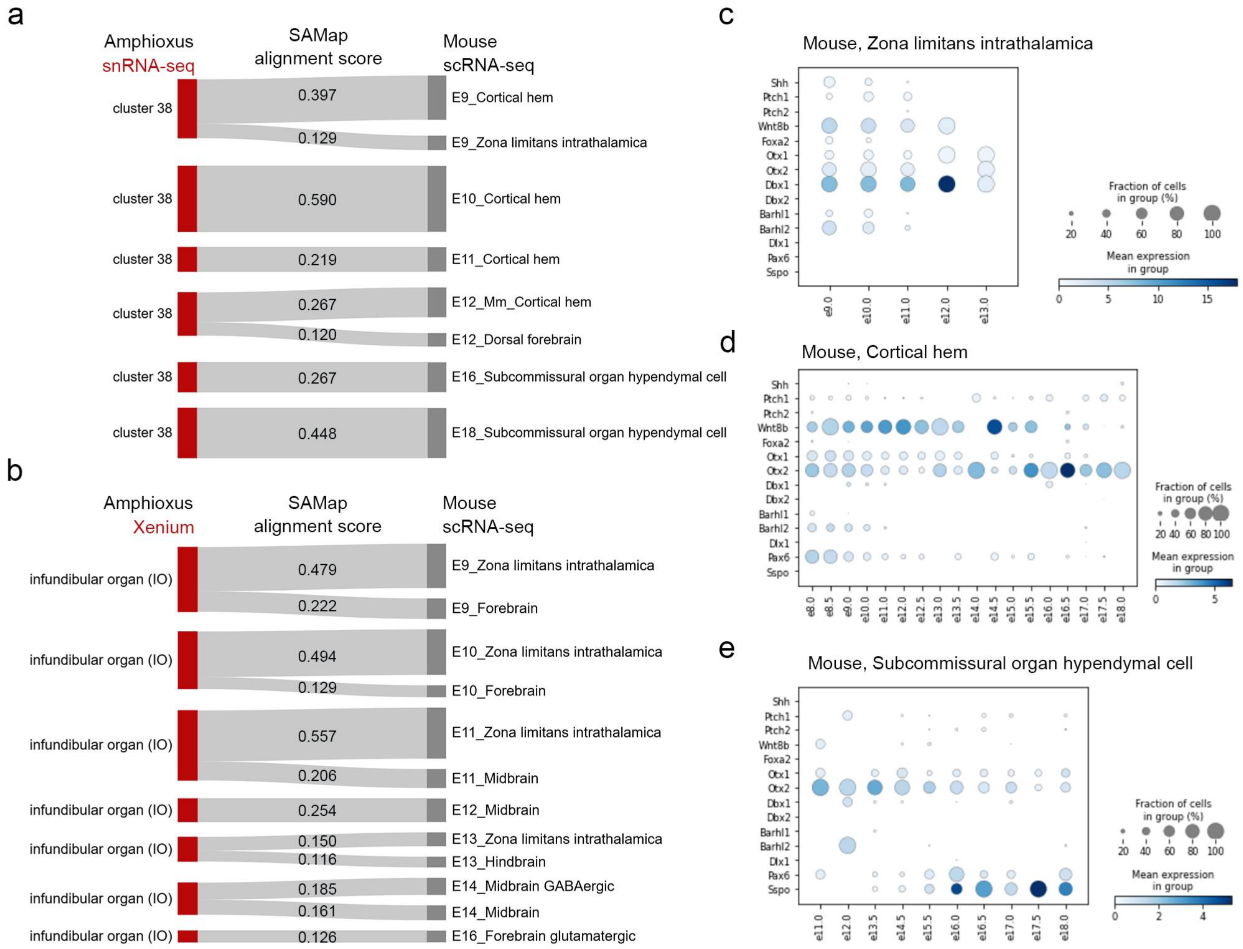


**Figure S18 | The amphioxus infundibular organ corresponds to three mouse structures: ZLI, cortical hem and subcommissural organ.**

**a-b**, Sankey plots showing SAMap-based cell-type alignments between cluster 38 of the amphioxus brain snRNA-seq dataset and the mouse brain dataset (**a**), as well as between the IO of the amphioxus Xenium dataset and the mouse brain dataset (**b**). Only alignments with scores > 0.1 are displayed. **c-e**, Dot plots of the mouse brain scRNA-seq data showing expression of ZLI-related genes within the ZLI (**c**), the cortical hem (**d**), and the subcommissural organ (**e**).

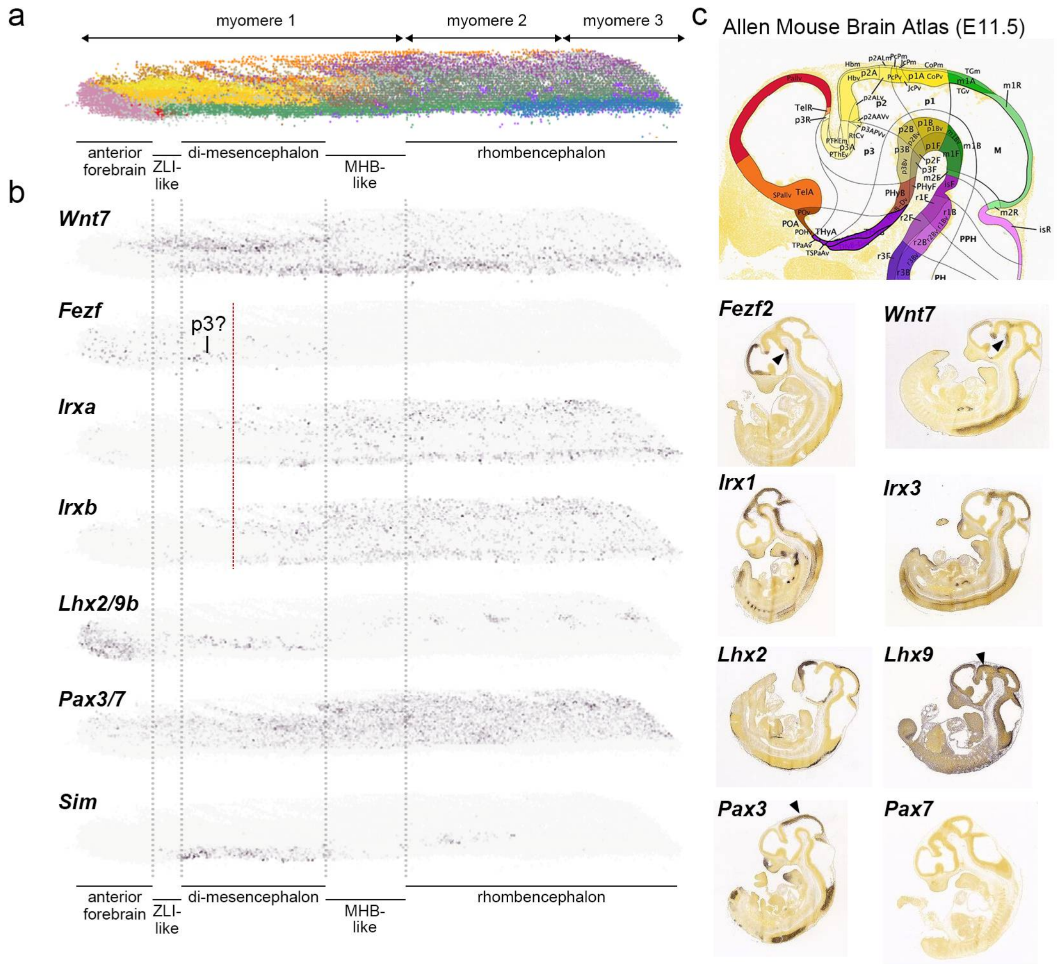


**Figure S19 | Expression profiles of genes associated with the vertebrate diencephalon.** **a**, Lateral view of the 3D point cloud model of the amphioxus brain, oriented with anterior to the left and colored by the anatomical brainregions. **b**, Lateral view of the 3D point cloud model showing the expression levels and domains of individual genes. Color scales were normalized for each gene, with the darkest tone representing the maximal expression. **c**, Expression profiles of representative diencephalic markers in E11.0 mouse embryos retrieved from the Allen Mouse Brain Atlas (*Fezf2* and *Wnt* as P3 markers; *Irx*a, *Irxb* and *Lhx9* as P2 marker; and *Pax3* as P2 and P1 marker).

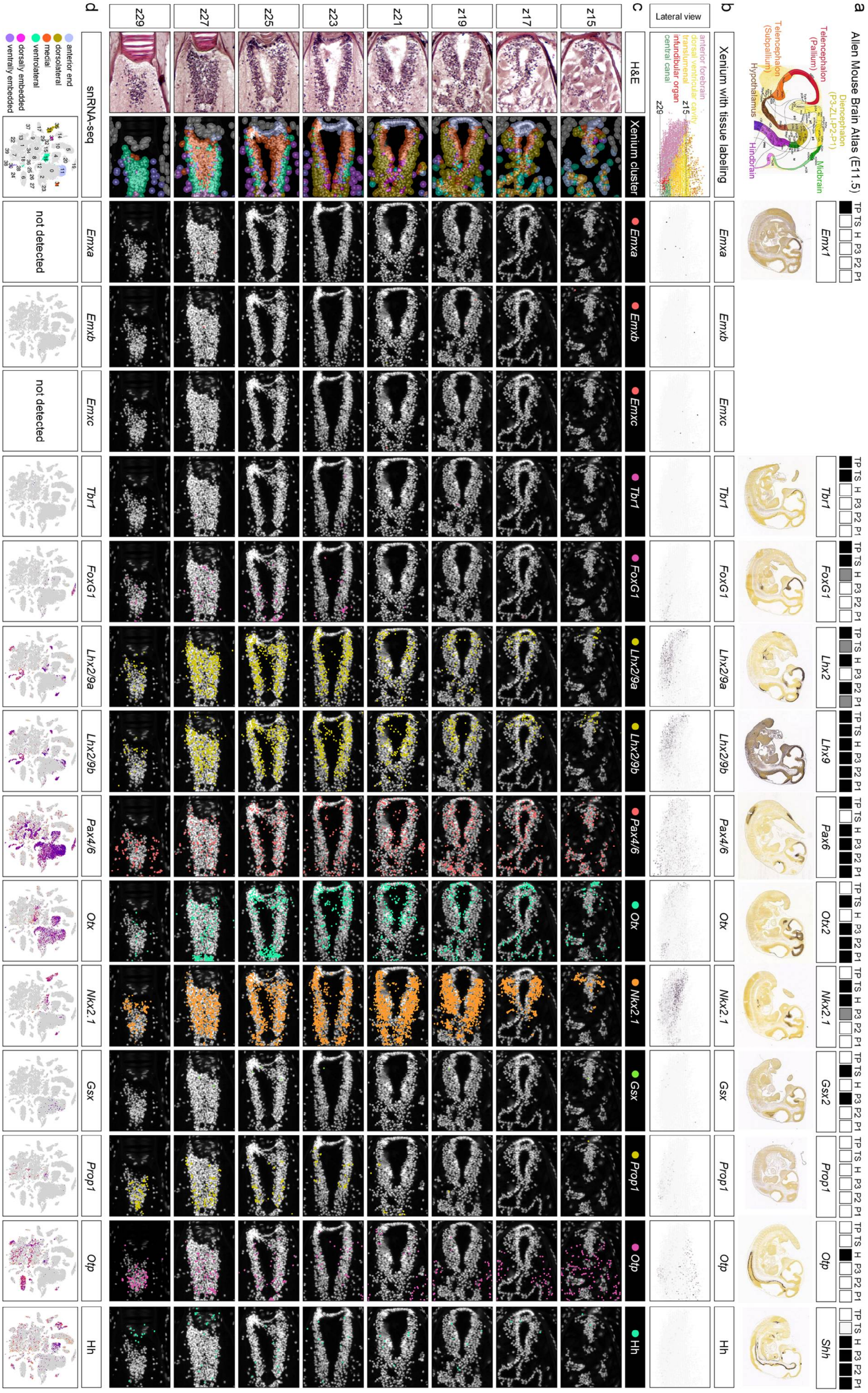


**Figure S20 | Expression profiles of genes associated with the vertebrate forebrain.**

**a**, Expression of the forebrain marker genes in E11.0 mouse embryos based on the Allen mouse brain atlas. **b**, Lateral view of the 3D point cloud model around the anterior forebrain of amphioxus, with cells colored by gene expression levels. Color scales were normalized for each gene, with the darkest tone representing the maximal expression. **c**, H&E-stained images and corresponding DAPI-stained Xenium spatial transcriptomic sections showing transcript localizations around the anterior forebrain. Each dot represents a single detected transcript. **d**, Projection of gene expression levels onto the t-SNE space of the snRNA-seq dataset.

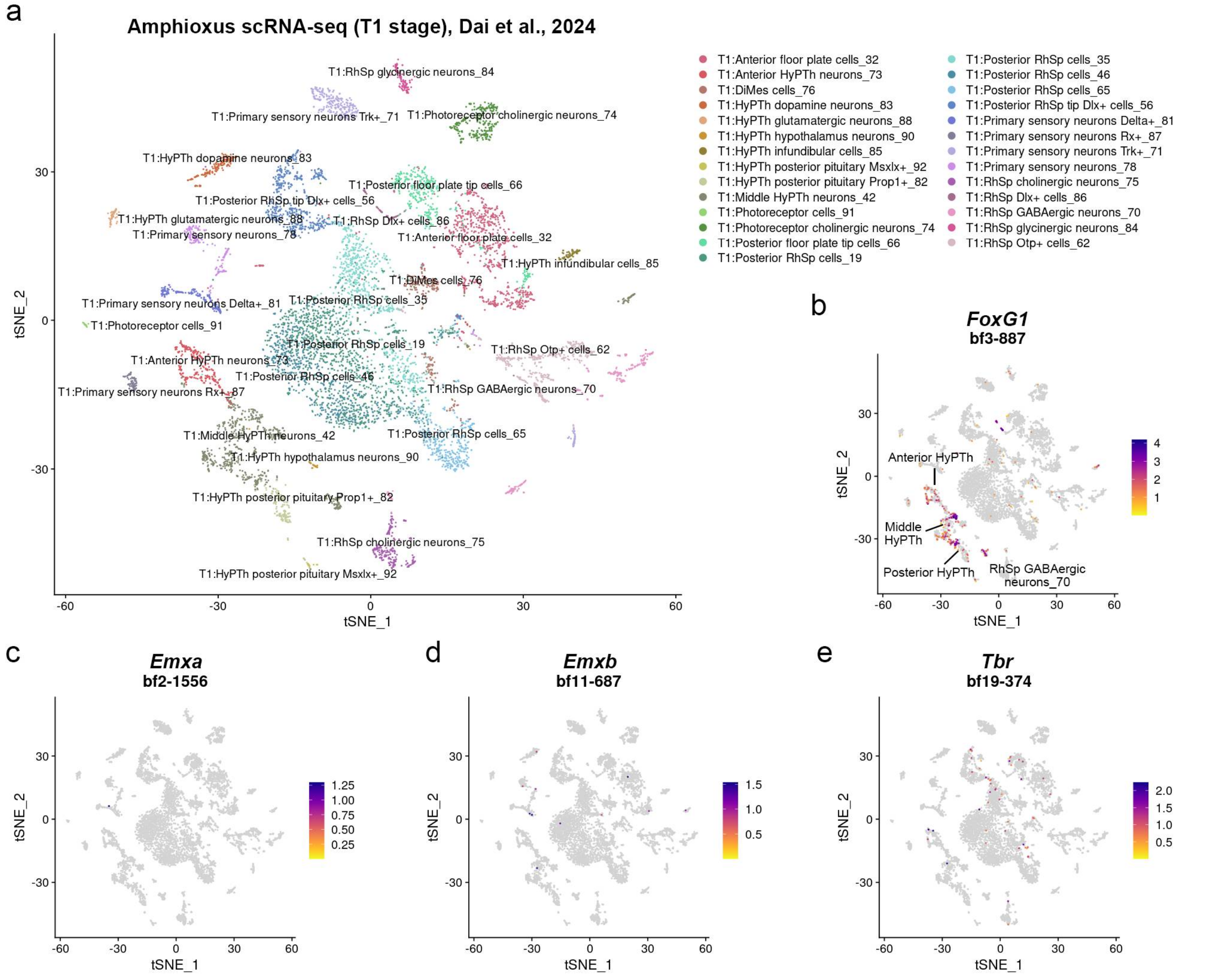


**Figure S21 | Expression of amphioxus orthologs of the vertebrate telencephalon genes in the amphioxus larval brain.**

**a**, t-SNE plot of the developing amphioxus nervous system annotated with cell types as described in the original publication[45]. **b-e**, Feature plots showing the expression of amphioxus orthologs of the vertebrate telencephalon marker genes, including *FoxG1* (**b**), *Emxa* (**c**), *Emxb* (**d**), and *Tbr* (**e**).

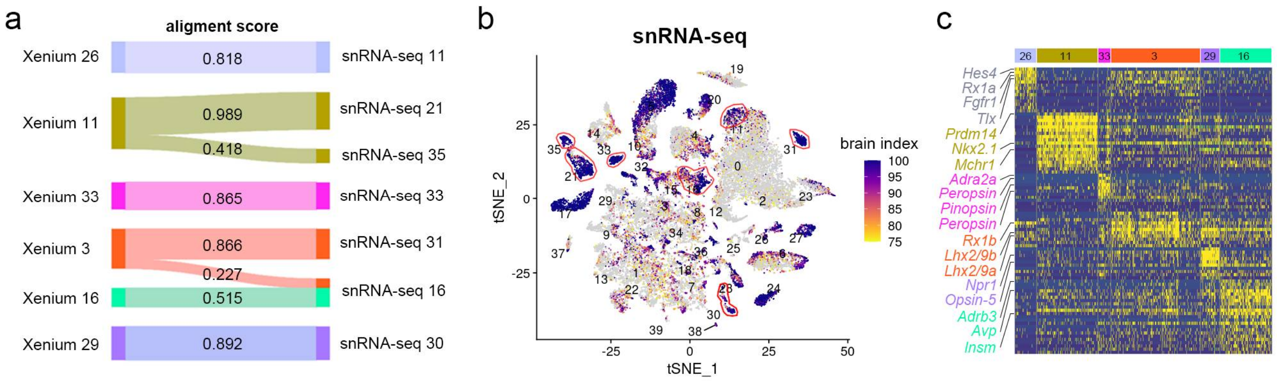


**Figure S22 | Anterior forebrain of the adult amphioxus brain.**

**a**, Sankey plot showing the correspondence between cell types in the Xenium and snRNA-seq datasets of the amphioxus brain. **b**, Projection of the brain-specific cell identity index onto the t-SNE space, with clusters corresponding to the anterior forebrain outlined in red. **c**, Heatmap showing the top differentially expressed genes from the Xenium dataset across anterior forebrain clusters.

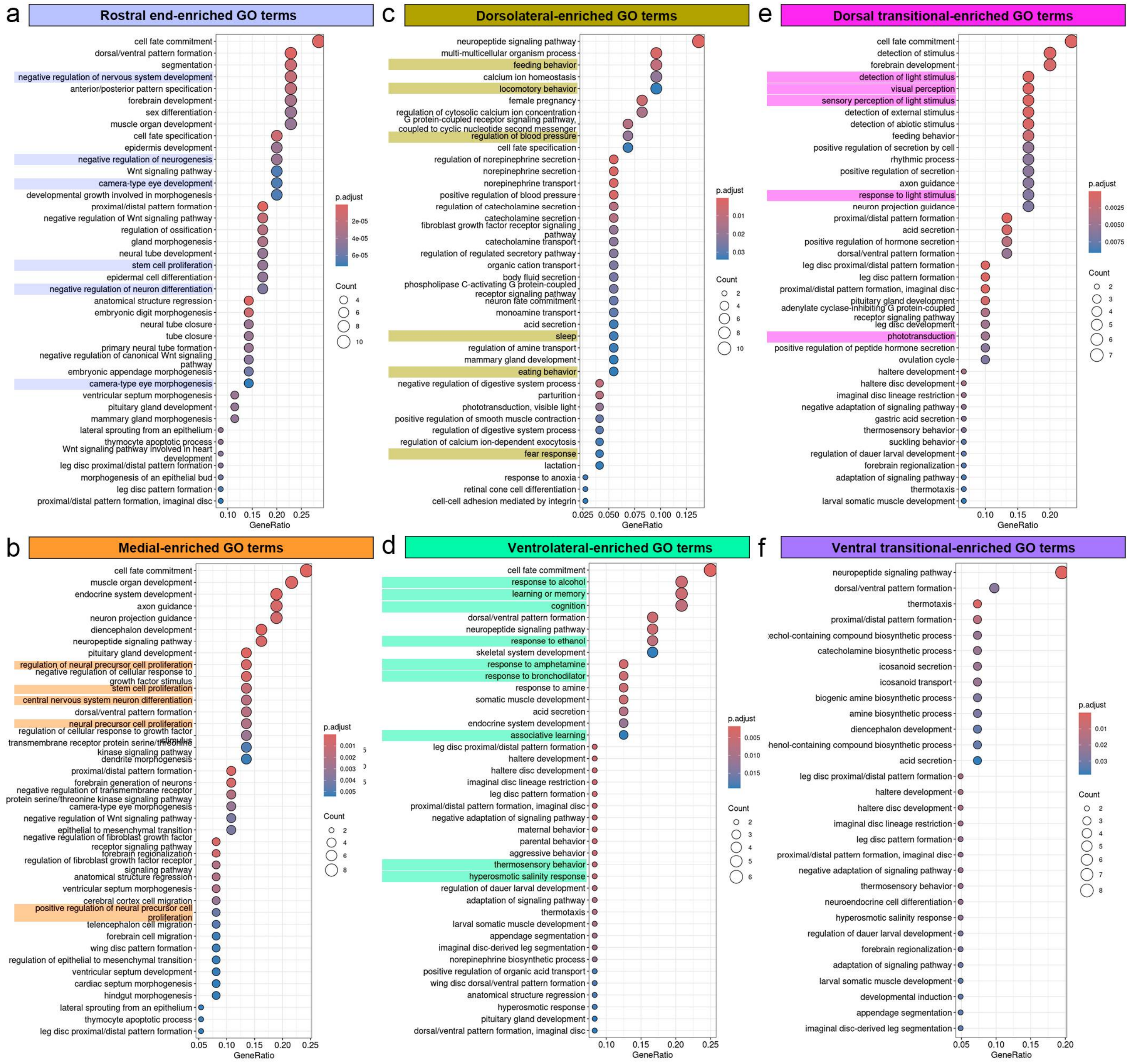


**Figure S23 | GO enrichment analyses of the anterior forebrain subregions.**

**a-f**, GO enrichment analyses of DEGs identified in subregions of the anterior forebrain, including rostral end (**a**), medial (**b**), dorsolateral (**c**), ventrolateral (**d**), dorsal transitional (**e**), and ventral transitional (**f**) regions. DEGs were derived from the snRNA-seq and Xenium datasets. The adjusted *p*-values < 0.05 are shown as colored circles. Circle size indicates the number of DEGs associated with each GO term.

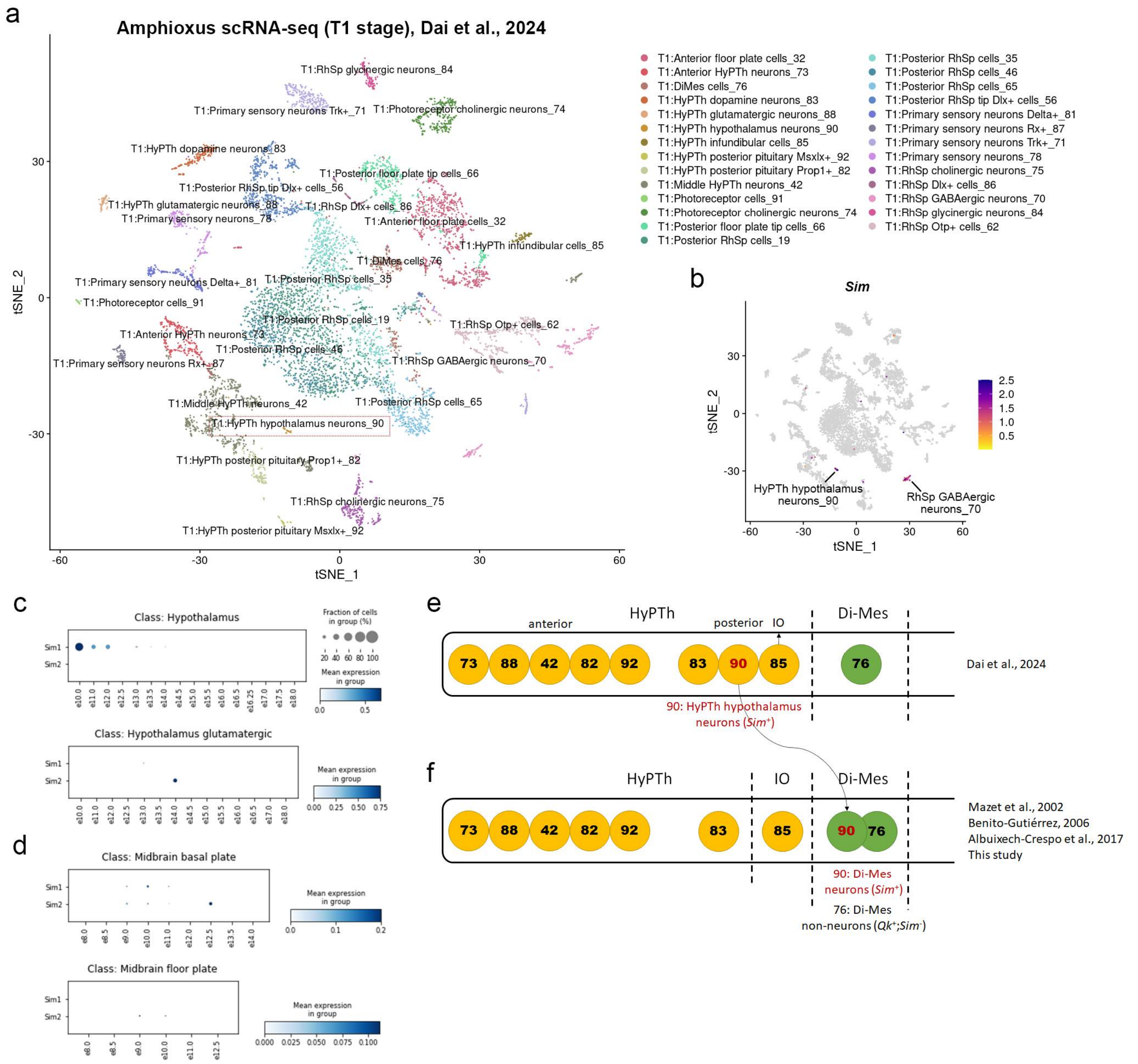


**Figure S24 | Identification of *Sim*$^{+}$ cells in the amphioxus brain.**

**a**, t-SNE plot of the developing amphioxus nervous system annotated with cell types as described in the original publication[45]. **b**, Feature plots showing the expression of the amphioxus *Sim* gene in the developing nervous system. **c-d**, Dot plots showing that mouse *Sim1* and *Sim2* genes are expressed in the hypothalamus (**c**) and midbrain (**d**), respectively. **e**, Previous scRNA-seq data suggested that the anterior *Sim* expression was localized to a hypothalamic region[45]. **f**, Our dataset reveals the anterior *Sim* expression predominantly in the Di–Mes region, consistent with several previous studies[17,58,59].
